# Determinants of telomere length across human tissues

**DOI:** 10.1101/793406

**Authors:** Kathryn Demanelis, Farzana Jasmine, Lin S. Chen, Meytal Chernoff, Lin Tong, Justin Shinkle, Mekala Sabarinathan, Hannah Lin, Eduardo Ramirez, Meritxell Oliva, Sarah Kim-Hellmuth, Barbara E. Stranger, Kristin G. Ardlie, François Aguet, Habibul Ahsan, GTEx Consortium, Jennifer Doherty, Muhammad G. Kibriya, Brandon L. Pierce

**Affiliations:** Department of Public Health Sciences, University of Chicago, Chicago, IL, USA; Section of Genetic Medicine, Department of Medicine, Institute for Genomics and Systems Biology, Center for Data Intensive Science, University of Chicago, Chicago, USA; New York Genome Center, New York, NY, USA; Statistical Genetics, Max Planck Institute of Psychiatry, Munich, Germany; Department of Systems Biology, Columbia University, New York, NY, USA; The Broad Institute of Massachusetts Institute of Technology and Harvard University, Cambridge, MA, USA; Department of Human Genetics, University of Chicago, Chicago, IL, USA; University of Chicago Comprehensive Cancer Center, Chicago, IL, USA; Department of Medicine, University of Chicago, Chicago, IL, USA; Huntsman Cancer Institute, University of Utah, Salt Lake City, UT, USA; The Broad Institute of MIT and Harvard, Cambridge, MA, USA; Cancer Center and Department of Pathology, Massachusetts General Hospital, Boston, MA, USA; Department of Genetics, Harvard Medical School, Boston, MA, USA; Program in Medical and Population Genetics, The Broad Institute of Massachusetts Institute of Technology and Harvard University, Cambridge, MA, USA; Stanley Center for Psychiatric Research, Broad Institute, Cambridge, MA, USA; Analytic and Translational Genetics Unit, Massachusetts General Hospital, Boston, MA, USA; Ocular Genomics Institute, Massachusetts Eye and Ear, Harvard Medical School, Boston, MA, USA; Department of Biomathematics, University of California, Los Angeles, Los Angeles, CA, USA; Section of Genetic Medicine, Department of Medicine, The University of Chicago, Chicago, IL, USA; Department of Biomedical Engineering, Johns Hopkins University, Baltimore, MD, USA; Department of Computer Science, Johns Hopkins University, Baltimore, MD, USA; Department of Genetic Medicine and Development, University of Geneva Medical School, Geneva, Switzerland; Population Health and Genomics, University of Dundee, Dundee, Scotland, UK; Department of Genetics, University of Pennsylvania, Perelman School of Medicine, Philadelphia, PA, USA; Department of Genetics, Washington University School of Medicine, St. Louis, Missouri, USA; Department of Pathology & Immunology, Washington University School of Medicine, St. Louis, Missouri, USA; Department of Genetics, Stanford University, Stanford, CA, USA; Division of Genetic Medicine, Department of Medicine, Vanderbilt University Medical Center, Nashville, TN, USA; Department of Biostatistics, University of Michigan, Ann Arbor, MI, USA; Institute for Genetics and Genomics in Geneva (iGE3), University of Geneva, Geneva, Switzerland; Swiss Institute of Bioinformatics, Geneva, Switzerland; Department of Computer Science, Princeton University, Princeton, NJ, USA; Center for Statistics and Machine Learning, Princeton University, Princeton, NJ, USA; Department of Computer Science, University of California, Los Angeles, Los Angeles, CA, USA; Program in Biomedical Informatics, Stanford University School of Medicine, Stanford, CA, USA; Department of Pathology, Stanford University, Stanford, CA, USA; Data Science Institute, Vanderbilt University, Nashville, TN, USA; Clare Hall, University of Cambridge, Cambridge, UK; MRC Epidemiology Unit, University of Cambridge, Cambridge, UK; Centre for Genomic Regulation (CRG), The Barcelona Institute for Science and Technology, Barcelona, Catalonia, Spain; Universitat Pompeu Fabra (UPF), Barcelona, Catalonia, Spain; Department of Epidemiology, Harvard T.H. Chan School of Public Health, Boston, MA, USA; Computer Science and Artificial Intelligence Laboratory, Massachusetts Institute of Technology, Cambridge, MA, USA; Department of Clinical Pharmacy, School of Pharmacy, University of Southern California, Los Angeles, CA, USA; Scripps Research Translational Institute, La Jolla, CA, USA; Department of Integrative Structural and Computational Biology, The Scripps Research Institute, La Jolla, CA, USA; Department of Statistics and Operations Research, Universitat Politècnica de Catalunya (UPC), Barcelona, Catalonia, Spain; Department of Statistics and Operations Research and Department of Biostatistics, University of North Carolina, Chapel Hill, NC, USA; Department of Public Health Sciences, The University of Chicago, Chicago, IL, USA; Department of Systems Pharmacology and Translational Therapeutics, University of Pennsylvania, Perelman School of Medicine, Philadelphia, PA, USA; Department of Genetics, Microbiology and Statistics, University of Barcelona, Barcelona. Spain; Departments of Biomedical Data Science and Statistics, Stanford University, Stanford, CA, USA; Department of Pathology and Laboratory Medicine, Ann & Robert H. Lurie Children’s Hospital of Chicago, Chicago, IL, USA; Center for Genetic Medicine, Department of Pharmacology, Northwestern University, Feinberg School of Medicine, Chicago, IL, USA; Department of Twin Research and Genetic Epidemiology, King’s College London, London, UK; Bioinformatics Research Center and Departments of Statistics and Biological Sciences, North Carolina State University, Raleigh, NC, USA; Department of Statistics, University of Chicago, Chicago, IL, USA; Department of Computer Sciences, Faculty of Sciences, University of Porto, Porto, Portugal; Instituto de Investigaça~o e Inovaça~o em Saúde, Universidade do Porto, Porto, Portugal; Institute of Molecular Pathology and Immunology, University of Porto, Porto, Portugal; Columbia University Mailman School of Public Health, New York, NY, USA; Life Sciences Department, Barcelona Supercomputing Center, Barcelona, Spain; Department of Clinical Biochemistry and Pharmacology, Ben-Gurion University of the Negev, Beer-Sheva, Israel; National Institute for Biotechnology in the Negev, Beer-Sheva, Israel; Leidos Biomedical, Frederick, MD, USA; Leidos Biomedical, Rockville, MD, USA; UNYTS, Buffalo, NY, USA; Washington Regional Transplant Community, Annandale, VA, USA; Therapeutics, Roswell Park Comprehensive Cancer Center, Buffalo, NY, USA; Gift of Life Donor Program, Philadelphia, PA, USA; LifeGift, Houston, TX, USA; Center for Organ Recovery and Education, Pittsburgh, PA, USA; LifeNet Health, Virginia Beach, VA. USA; National Disease Research Interchange, Philadelphia, PA, USA; Van Andel Research Institute, Grand Rapids, MI, USA; Department of Neurology, University of Miami Miller School of Medicine, Miami, FL, USA; Biorepositories and Biospecimen Research Branch, Division of Cancer Treatment and Diagnosis, National Cancer Institute, Bethesda, MD, USA; Temple University, Philadelphia, PA, USA; Virgina Commonwealth University, Richmond, VA, USA; European Molecular Biology Laboratory, European Bioinformatics Institute, Hinxton, United Kingdom; Genomics Institute, UC Santa Cruz, Santa Cruz, CA, USA; Carl Icahn Laboratory, Princeton University, Princeton, NJ, USA; Department of Population Health Sciences, The University of Utah, Salt Lake City, Utah, USA; Schools of Medicine, Engineering, and Public Health, Johns Hopkins University, Baltimore, MD, USA; Department of Biostatistics, Bloomberg School of Public Health, Johns Hopkins University, Baltimore, MD, USA; Department of Medical Biology, The Walter and Eliza Hall Institute of Medical Research, Parkville, Victoria, Australia; Altius Institute for Biomedical Sciences, Seattle, WA, USA; Division of Genetics, University of Washington, Seattle, WA, University of Washington, Seattle, WA, USA; Department of Cardiology, University of Washington, Seattle, WA, USA; HudsonAlpha Institute for Biotechnology, Huntsville, AL, USA; Genome Sciences, University of Washington, Seattle, WA, USA; National Institute of Dental and Craniofacial Research, Bethesda, MD, USA; Division of Neuroscience and Basic Behavioral Science, National Institute of Mental Health, National Institutes of Health, Bethesda, MD, USA; National Institute on Drug Abuse, Bethesda, MD, USA; Office of Strategic Coordination, Division of Program Coordination, Planning and Strategic Initiatives, Office of the Director, National Institutes of Health, Rockville, MD, USA; Division of Genomic Medicine, National Human Genome Research Institute, Bethesda, MD, USA

**Author notes:** Corresponding Author. (B.L.P).

## Abstract

Telomere shortening is a hallmark of aging. Telomere length (TL) in blood cells has been studied extensively as a biomarker of human aging and disease; however, little is known regarding variability in TL in non-blood, disease-relevant tissue types. Here we characterize variability in TL measurements for 6,391 tissue samples, representing >20 tissue types and 952 individuals from the Genotype-Tissue Expression (GTEx) Project. We describe differences across tissue types, positive correlation among tissue types, and associations with age and ancestry. We show that genetic variation impacts TL in multiple tissue types, and that TL can mediate the effect of age on gene expression. Our results provide the foundational knowledge regarding TL in healthy tissues that is needed to interpret epidemiological studies of TL and human health.

**ONE SENTENCE SUMMARY:** Telomere length varies by tissue type but is generally correlated among tissue types (positively) and with age (negatively).

## MAIN TEXT

Telomeres are DNA-protein complexes located at the end of chromosomes that protect chromosome ends from degradation and fusion (*1*). The length of the DNA component of telomeres shortens as cells divide (*2*) with short telomeres eventually triggering cellular senescence (*3, 4*). In most human tissues, TL gradually shortens over the life course, and TL shortening is considered a hallmark (and a potential underlying cause) of human aging (*5*). In human studies, short TL measured in leukocytes is associated with increased risk of aging-related diseases including cardiovascular disease (*6*) and type II diabetes (*7*) as well as all-cause mortality (*8*). However, long TL may increase risk for some types of cancer (*9-11*). Leukocyte TL is influenced by inherited genetic variation (single nucleotide polymorphisms [SNPs]), some of which reside near genes with roles in telomere maintenance (*12-15*). Leukocyte TL is also associated with lifestyle factors (e.g., obesity) and exposures (e.g., cigarette smoking) (*16, 17*).

Epidemiologic studies of TL predominantly use blood (occasionally saliva) as a DNA source. Thus, our understanding of variation in TL, its determinants (e.g., demographic, lifestyle, and genetic factors), and its associations with disease phenotypes is based almost entirely on TL measured in leukocytes from whole blood. Few prior studies have compared TL in leukocytes to TL in other human tissue types; these prior studies are relatively small (<100 participants; <5 tissue types) but provide evidence that TL differs across tissue types and that TL measurements from different tissue types are correlated (*18, 19*). However, larger studies of many additional tissue types are needed to gain a comprehensive understanding of variation in TL and its determinants within and across a wide range of human tissues and cell types. In order to address these gaps in our understanding of TL and its role as a biomarker of aging and disease risk, we measured TL in > 6,000 unique tissue samples, representing >20 distinct tissue types and > 950 individual donors from the Genotype-Tissue Expression (GTEx) Project (see **Methods**) (*20*). In this paper we (1) characterize sources of variation in TL, (2) evaluate leukocyte TL as a proxy for TL in other disease-relevant tissues, (3) describe the relationship between age and TL across tissue types, and (4) describe biological determinants and correlates of TL.

We attempted measurement of relative TL (telomere repeat abundance relative to a standard/reference DNA sample [RTL]) for 7,234 tissue samples from 962 GTEx donors using a Luminex-based assay [see **Methods**]. After removing 836 samples with failed RTL measurements and seven RTL measures that were within-tissue outliers, our analytic dataset included 6,391 tissue-specific RTL measurements from 952 donors, with 24 different tissue types having ≥ 25 RTL measurements (**Table S1**). On average, each donor had RTL measured in seven different tissue types (range: 1-26 tissue types) (**Figure S1**). The median donor age was 55 (range: 20-70) years, there were more males (67%) than females, and participants were primarily white (85%) (**Table S1**).

### TL varies across (and correlates among) human tissues types

We estimated the contribution of tissue type to the variation in RTL using linear mixed models (LMMs) that were adjusted for fixed effect covariates (age, sex, BMI, race/ethnicity, donor ischemic time, and technical factors [DNA concentration and sample plate]) and random effects of tissue type and donor (see **Methods** and **Table S2**). On average, RTL was the shortest in whole blood (WB) and longest in testis, with testis being a clear outlier (*p* < 2×10^−16^ compared to all other tissues) (**Figure 1A**). Tissue type explained 24.3% of the variation in RTL across all tissues but only 11.5% when testis was excluded. We examined pairwise correlations (Pearson) in RTL among tissue types with tissue pairs from same donor, restricting to 20 tissue types with TL data for ≥ 75 samples (**Figure 1B**). Forty tissue-pair correlations passed a Bonferroni *p*-value threshold (*p* < 3 × 10^−4^) and all correlations were positive (**Data Table S1**). All tissue pairs from the same organ were among the stronger correlations observed: sun exposed and non-exposed skin (r=0.24, *p=*9 × 10^−3^, n=112), transverse and sigmoid colon (r=0.40, *p=*8 × 10^−7^, n=139), and esophagus mucosa (EM) and gastric junction (EGJ) (r=0.22, *p=*3 × 10^−3^, n=188). Applying hierarchical clustering to these pairwise correlations using average linkage, tissue RTL separated into three clusters (**Figure 1B** and **Figure S2**). Two clusters were characterized by common developmental origin: 1) mesodermal and ectodermal (e.g., muscle and skin) and 2) endodermal origin tissues (e.g., stomach and lung). Thyroid and brain cerebellum formed the third cluster. Similar clustering patterns among tissue types were observed for females (**Figure S3**) and males (**Figure S4**), where testis was also an outlying tissue type and clustered with thyroid. The positive correlations observed among most tissue types are most likely due to the fact that the initial TL in the zygote impacts TL in all adult tissues through mitotic inheritance; while differences among tissue types are likely attributable to variability in both intrinsic (e.g., cell division rate/history, telomere maintenance) and extrinsic (e.g., response to environmental exposures) factors across tissues (**Figure 1D**).

**Figure 1.**
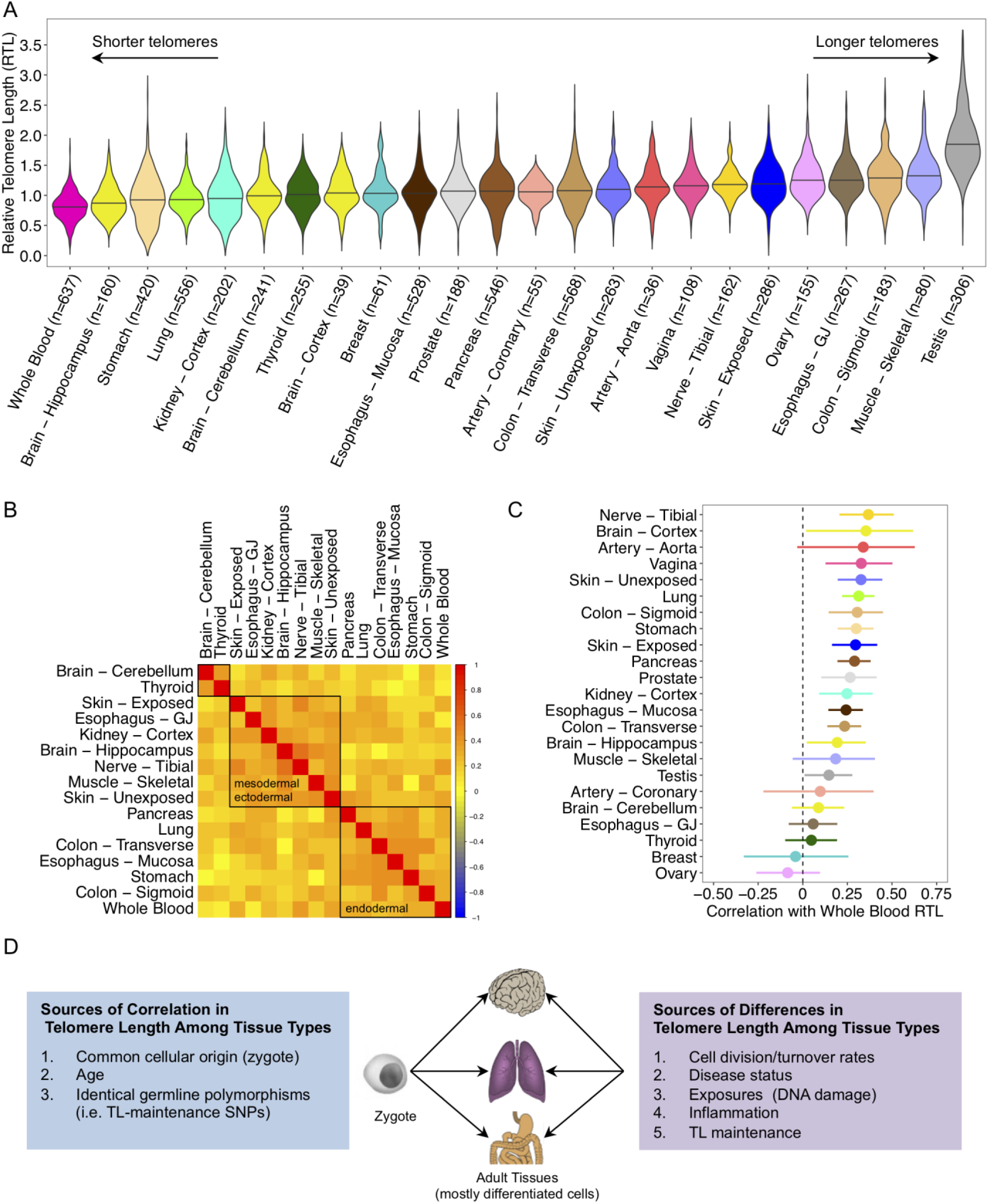
Telomere lengths differ across human tissue types but are correlated among tissues types. A) Distribution of RTL across 24 GTEx tissue types (ordered by median RTL). B) Pearson (r) correlations between RTL measures from different tissue types. Tissues included have ≥ 75 samples and were not sex-specific. Red, yellow, and blue correspond r=1, r=0, and r=−1, respectively. Black boxes are results from hierarchical clustering for k=3 clusters (exact correlations are in **Data Table S1**). C) Pearson correlations between whole blood RTL and tissue-specific RTL measures (with 95% confidence intervals). D) Theoretical framework describing determinants of telomere length across human tissue types.

### Whole blood TL is a proxy for TL in other tissue

WB RTL was positively correlated (*p* < 0.05) with tissue-specific RTL measures from 15 out of 23 tissue types (each with n ≥ 25), with Pearson correlations ranging from 0.15 to 0.37 (**Figure 1C**). These results demonstrate that WB RTL can serve as a proxy for TL in many tissue types. WB RTL captured between 2% (testis) and 14% (tibial nerve) of the variation in RTL measured in other tissue types. Adjustment for age, sex, body mass index (BMI), and donor ischemic time did not have a major impact the associations observed between WB RTL and tissue RTL for our 23 tissue types (**Figure S5**).

RTL measures have inherent measurement error (*21*), including our Luminex assay (*22*), so we simulated data to determine the extent to which measurement error (i.e., random, non-differential error) is expected to impact the observed correlation (r) estimates. We simulated data representing TL measures from two tissue types, and varied the value of the true Pearson correlation (r) between TL measures from those tissues and the amount of error present in the RTL measurement (see **Methods**). An inverse linear relationship was observed between the proportion of variation that measurement error accounted for in the RTL measure and the observed r value (**Figure S6**). When measurement error accounted for 50% of the variation in both RTL measurements (most extreme scenario tested), the observed r between the two tissues decreased by ∼50% compared to the true r, and the correlation between an error-prone measure and the true TL (in another tissue) decreased by ∼25%. Thus, the correlations we observe between WB RTL and RTL in other tissue types are likely underestimates of the true correlations.

Previously we have shown the Luminex-based TL method to have a correlation (r) of ∼0.7 (a r^2^ of ∼0.5) with the Ssouthern blot analysis of terminal restriction fragment lengths method for TL measurement (in a blinded comparison study) (*22*). The Southern blot method is a more expensive and labor-intensive approach for TL measurement with much higher DNA input requirements. If we consider Southern blot to be the gold standard for TL measurement, thereby assuming that 50% of the variation in our Luminex-based measure is random error, then the correlations observed in this study are ∼50% lower than the true correlations (based on **Figure S6**).

### TL varies among individuals and by participant characteristics

TL varied across individuals (donors) (**Figure 2A** [top]), explaining 8.7% of variation in RTL across all tissues and 11.2% with testis excluded (based on estimates from adjusted LMM) (**Table S2**). Adjusting for tissue type and donor (as random effects), age explained 3.3% (among all tissues) and 4.4% (excluding testis) of variation in RTL while BMI, TL-associated SNPs, and race/ethnicity each explained less than 1% of the variation across all tissues (marginal R^2^, *p* < 0.001) (**Figure 2B** [top]). We observed no association between sex and RTL across all tissues (**Table S2**), and sex showed very little evidence of association with RTL in tissue-specific analyses (**Table S3**). We conducted a principal component (PC) analysis of RTL from eleven non-reproductive tissue types (each with n ≥ 200 samples) from 750 participants (see **Methods**) and generated a composite measure of TL based on the first PC that explains 51% of the variation in TL among these tissue types **(Figure 2A** [bottom]). We observed that age and BMI were associated with shorter composite RTL and explained 13.7% and 1.3%, respectively, of the variation in this composite TL measure (**Figure 2B** [bottom]). Race/ethnicity was associated with longer composite TL in African Americans compared to white individuals and explained 1.6% of the variation in composite TL. This composite TL likely reflects the TL in the zygote (and in tissues during early development) that is mitotically inherited by cells in adult tissues.

**Figure 2.**
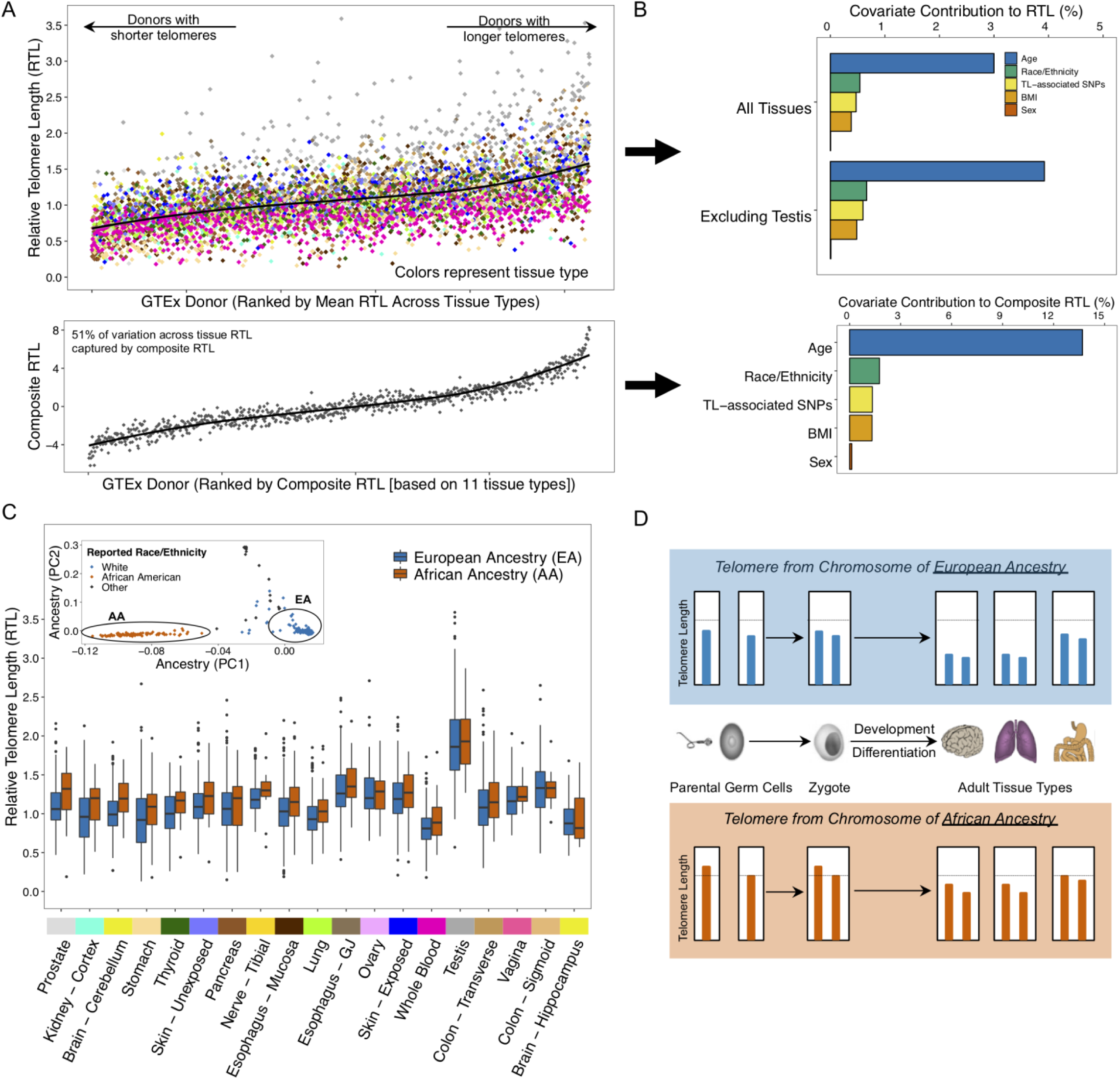
Telomere length varies among individuals and by ancestry. A) Distribution of RTL across GTEx donors ranked by donors’ mean RTL across all measured tissue types (top panel). Bottom panel shows distribution of a “composite RTL” measure, estimated based on the first principal component from an analysis of 11 tissue types (see **Methods**). Colors correspond to GTEx tissue type. B) Contribution of selected covariates to variability in RTL across all tissues (top) and composite RTL (bottom). For the analysis across all tissues, estimates were extracted as marginal R^2^ values from linear mixed models adjusted for tissue type and donor as random effects. C) Distribution of RTL measures for individuals of European and African ancestry. Tissue types are ranked by largest difference between median RTL between the two ancestry groups. Interior panel shows genotyping principal components (PCs), demonstrating consistent clustering of individuals by genetically predicted ancestry. Associations between African ancestry and RTL are presented in **Table S4**. D) Schematic describing the direct inheritance of TL from parental germ cells and expected relationship to TL across adult tissue types for individuals of African and European ancestry. Genetic (and reported race/ethnicity) ancestry was color coded for African (red) and European (blue) in panels C and D.

### TL is longer in genomes of African Ancestry

To further explore differences in TL by race/ethnicity, we first confirmed that PCs derived from genome-wide SNP data (n=831 donors), representing genetic ancestry, showed clear clustering by reported race/ethnicity among donors (**Figure 2C** [upper left]). Genetic ancestry (European vs. African) explained 0.6% of the variation in RTL across all tissues (marginal R^2^, *p=*1×10^−5^) after adjusting for tissue type and donor as random effects and 2.3% of the variation in the composite RTL measure (*p=*7×10^−5^). After including adjustments for age, sex, donor ischemic time, technical factors (DNA concentration and sample plate) and random effects of tissue type and donor, RTL was longer among individuals of African ancestry compared to individuals of European ancestry in analyses of all tissue types combined (*p=*0.007), consistent with prior studies of leukocyte TL (*23-26*). The adjusted association between African ancestry and RTL was positive for 17 out of 20 tissues tested, with *p*-values < 0.05 for brain cerebellum (*p=*0.03), thyroid (*p=*0.02), prostate (*p=*0.03), lung (*p=*0.02), and whole blood (*p=*0.005) (**Figure 2C** and **Table S4**). The observation that individuals of African ancestry have longer TL in many tissue types is consistent with the hypothesis that ancestry-based differences in TL are present early in development (*27*) and potentially in germ cells (pre-conception). In other words, our results suggest that offspring (zygotes) inherit telomeres from germ cells that vary in TL due to ancestry, and these ancestry-based differences in TL are mitotically transmitted to daughter cells, and eventually to cells in many adult tissue types. This “direct transmission” of TL from parent to offspring (*28*) would result in the observed ancestry-based differences across many tissue types (summarized in **Figure 2D**). One likely cause of this ancestry-based difference is natural section on SNPs know to impact TL (*29*), although selection on TL itself could also contribute.

### TL is correlated with age in most tissues

Of 24 tissues with ≥ 25 samples, RTL was negatively correlated (r < 0) with age in 21 tissue types (*p* < 0.05 in 14 tissue types) (**Figure 3A, Figure S7**), supporting the hypothesis that age-related TL shortening occurs in most tissue types. The strongest correlations with age were observed for WB (r=−0.35, *p=*2×10^−19^, n=637) and stomach (r=−0.37, *p=*7×10^−15^, n=420) (**Table S5**). Age explained more of the variation in RTL for tissues with shorter mean RTL (r^2^=0.23, *p=*0.02) (**Figure 3B**). Among tissue types in which RTLs did not have a clear correlation with age (*p* > 0.05), we examined whether RTL differed among 5-year age groups, but we observed no differences in RTL among 5-year age groups for testis, ovary, cerebellum, vagina, skeletal muscle, thyroid, and EGJ. While prior studies have observed longer TL in sperm from older men (*30*), we did not observe a clear increasing (or decreasing) trend for testis RTL with increasing age (**Figure S8**).

**Figure 3.**
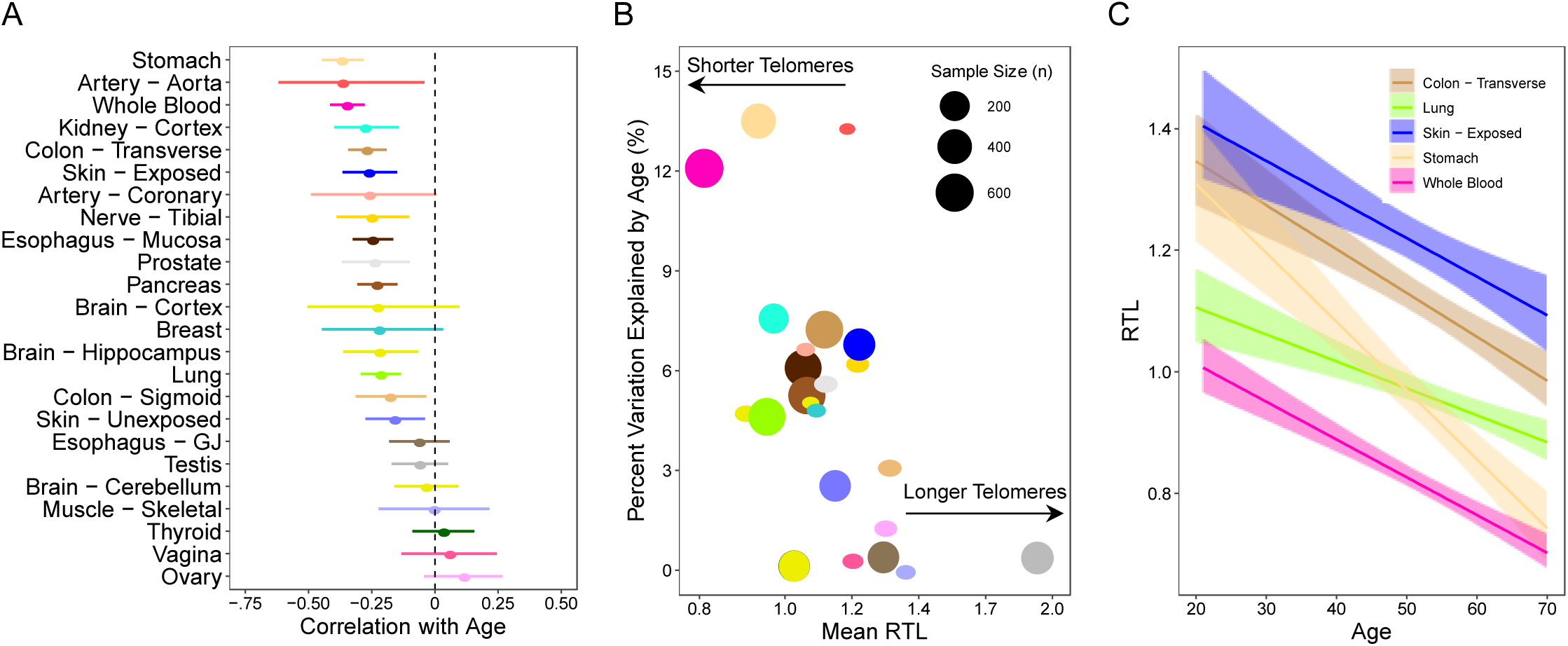
Age is negatively correlated with telomere length in most tissues, and correlation is strongest in tissues with shorter telomeres. A) Pearson correlations between age and tissue-specific RTL measures. B) Scatterplot of mean RTL for each tissue versus the percent variation explained by age (r^2^) for each tissue. The size of each point is proportional to sample size for that tissue type. C) Relationship between RTL and age for five selected tissue types (whole blood, lung, stomach, transverse colon, and skin [exposed]). For all plots, colors correspond to tissue type.

Among tissue types for which RTL was associated with age (*p* < 0.05), the strength of association varied across tissue types (**Figure 3C** and **Table S5**). To further explore the hypothesis that TL shortens at different rates in different tissue types, we calculated the difference in RTL (ΔRTL) between all pairs of tissue types available for each donor. We constructed 155 ΔRTL variables restricting to tissues pairs with complete data for ≥ 50 donors. The Pearson correlation between ΔRTL and age was estimated for each tissue type pair to determine if the ΔRTL varies with age (**Figure S9**). Forty-two of the 155 ΔRTL variables were correlated with age (*p* < 0.05), and the absolute values of these correlations ranged from 0.12 to 0.38 (**Data Table S2**). Four of the ΔRTLs surpassed Bonferroni *p*-value of 3 × 10^−4^: EGJ and stomach (r=0.32, *p=*1 × 10^−5^, n=176), WB and thyroid (r=0.30, *p=*3 × 10^−5^, n=182), EM and stomach (r=0.25, *p=*3 × 10^−5^, and n=276), and WB and ovary (r=0.33, *p=*2 × 10^−4^, n=120). Our results indicate that age explains up to 14% of the variation in the difference in RTL between pairs of tissue types. A prior study of 87 adults reported that the age rate of TL shortening was similar for muscle, leukocytes, fat, and skin (i.e. no association between age and ΔRTLs), concluding that age-related TL loss within stem cells is consistent across adult tissue types (*18*). When we examined these tissue types among our ΔRTL pairs (n ≥ 50), age was correlated with ΔRTL for skeletal muscle and blood (r=0.37, *p=*2×10^−3^, n=68) but less for skin (unexposed) and blood (r=0.09, *p=*0.2, n=197) and skin (exposed) and blood (r=0.08, *p=*0.24, n=200).

### Leukocyte TL-associated genetic variants and TL in other tissues

Prior genome-wide association studies (GWAS) have identified SNPs associated with leukocyte TL (*12-15*). We constructed a weighted polygenic SNP score for each donor using nine leukocyte TL-associated SNPs, with higher score reflecting longer TL (see **Methods** and **Table S6**) (*31*). We examined the association between this polygenic SNP score and RTL for tissue types with ≥ 100 samples. After adjustment for age, sex, genotyping PCs, donor ischemic time, and technical factors (DNA concentration and sample plate) as a random effect, an association with the SNP score (*p* < 0.05) was observed for WB RTL (*p=*0.007) (**Figure S10**), cerebellum RTL (*p=*0.03), pancreas RTL (*p=*0.04), and transverse colon RTL (*p=*0.02) (**Figure 4A, Figure S11**, and **Table S7**). Among all 18 tissue types, 16 had positive association estimates (binomial test [p_0_=0.5], *p=*0.001). In analyses of all tissue types, RTL was positively associated with this SNP score (*p=*0.01) after adjustment for age, sex, genotyping PCs, donor ischemic time, and technical factors (DNA concentration and sample plate) and random effects of tissue type and donor. These results indicate that at least some of the genetic variants (or regions) that impact leukocyte TL also impact TL in other tissue types.

**Figure 4.**
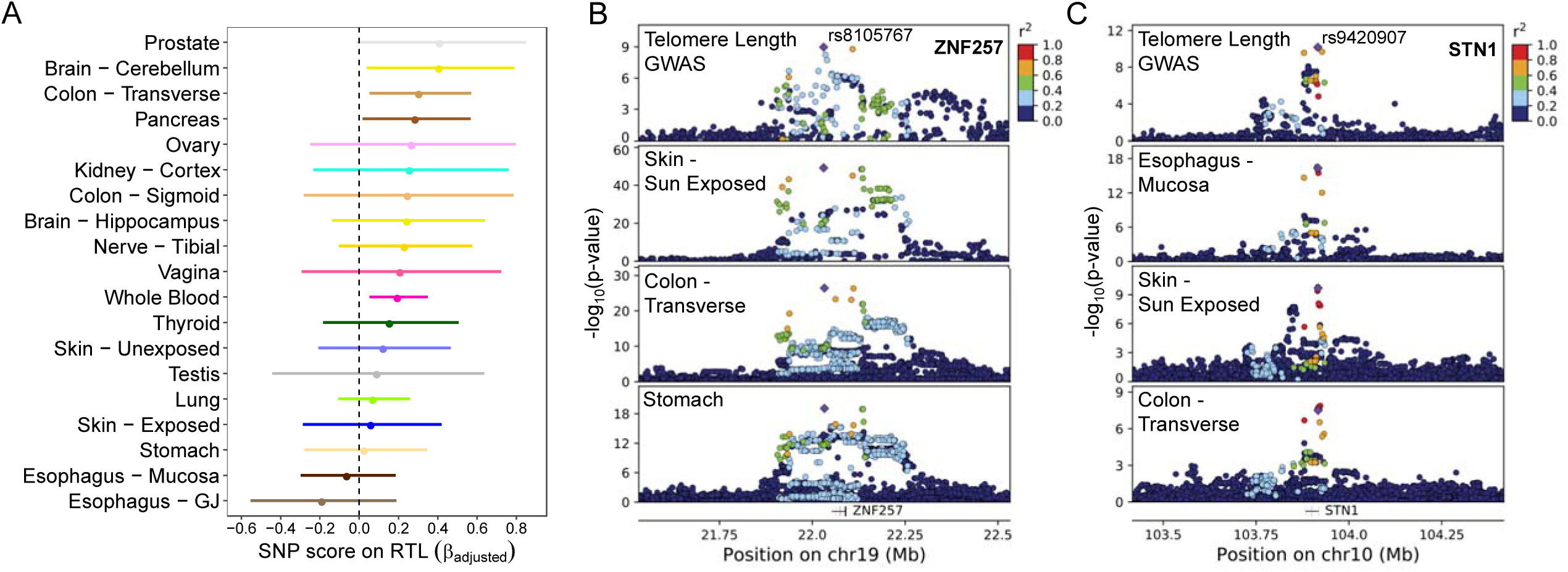
Genetic determinants of leukocyte telomere length impact telomere length in other tissue types and expression of nearby genes. A) Associations between a polygenic SNP score for leukocyte TL and tissue-specific RTL measures. Colors correspond to tissue type. B) Leukocyte TL association signal co-localizes with a *cis*-eQTL (expression quantitative trait locus) for *ZNF257* (∼40 kb upstream of *ZNF208*). Top plot shows results from the ENGAGE consortium GWAS of leukocyte TL, and bottom three plots correspond to *cis*-eQTL results from GTEx tissues: skin sun exposed, colon – transverse, and stomach. C) Leukocyte TL association signal co-localizes with a *cis*-eQTL for *STN1* (a.k.a., *OBFC1* in hg19). Top plot corresponds to results from the ENGAGE consortium GWAS of leukocyte TL, and bottom three plots correspond to *cis*-eQTL results from GTEx tissues: skin sun exposed, esophagus – mucosa (EM), and colon – transverse.

### TL-associated variants influence local gene expression

Among the nine regions known to harbor SNPs associated with leukocyte TL, we examined whether these loci also affected local gene expression in GTEx tissue types and cell lines (see **Methods**). We used co-localization analysis to estimate the probability that a common causal variant underlies association signals for leukocyte TL (from GWAS) (*12-14*) and *cis*-eQTL (expression quantitative trait loci) association signals from GTEx (v8) analyses (GTEx Consortium 2019 [GTEx main paper]). Co-localization results indicated that at least six of the nine TL-associated regions shared a common causal variant with a *cis*-eQTL in at least one tissue type, based on a posterior probability of co-localization of ≥ 80% across all three sets of priors tested (see **Methods**) (**Figure 4B, Figure 4C, Figure S12**, and **Data Table S3**). The association signal for TL on chromosome 19 (represented by rs8105767) showed strong evidence of co-localization with an eQTL affecting expression of *ZNF257* in eight tissue types, including skin (sun exposed), transverse colon, and stomach (**Figure 4B**). The association signal for TL on chromosome 10 (represented by rs9420907) co-localized with an eQTL affecting expression of *STN1* in seven tissue types, including skin (sun exposed), transverse colon, and EM (**Figure 4C**). Additional TL-associated loci showed co-localization with GTEx eQTLs for *NAF1, MYNN, RP11-109N23*.*6*, and *TSPYL6* (**Figure S12** and **Data Table S3**). These results suggest that TL-associated loci influence TL within human tissues via regulation of the expression of genes known to be involved in telomere maintenance (e.g., *STN1, NAF1*) (*12*), as well as genes whose role in telomere maintenance is unclear (e.g., *ZNF257*). Notably, we observed very little evidence of co-localization of the *TERT* or *TERC* TL-associated regions with any *cis*-eQTLs, likely because *TERT* and *TERC* have low or undetectable expression in a majority of adult GTEx tissue samples (**Figure 5A**). This suggests that eQTL studies of cells from stem and/or developmental tissues may be needed to understand the mechanisms underlying genetic regulation of *TERT* and *TERC* expression.

**Figure 5.**
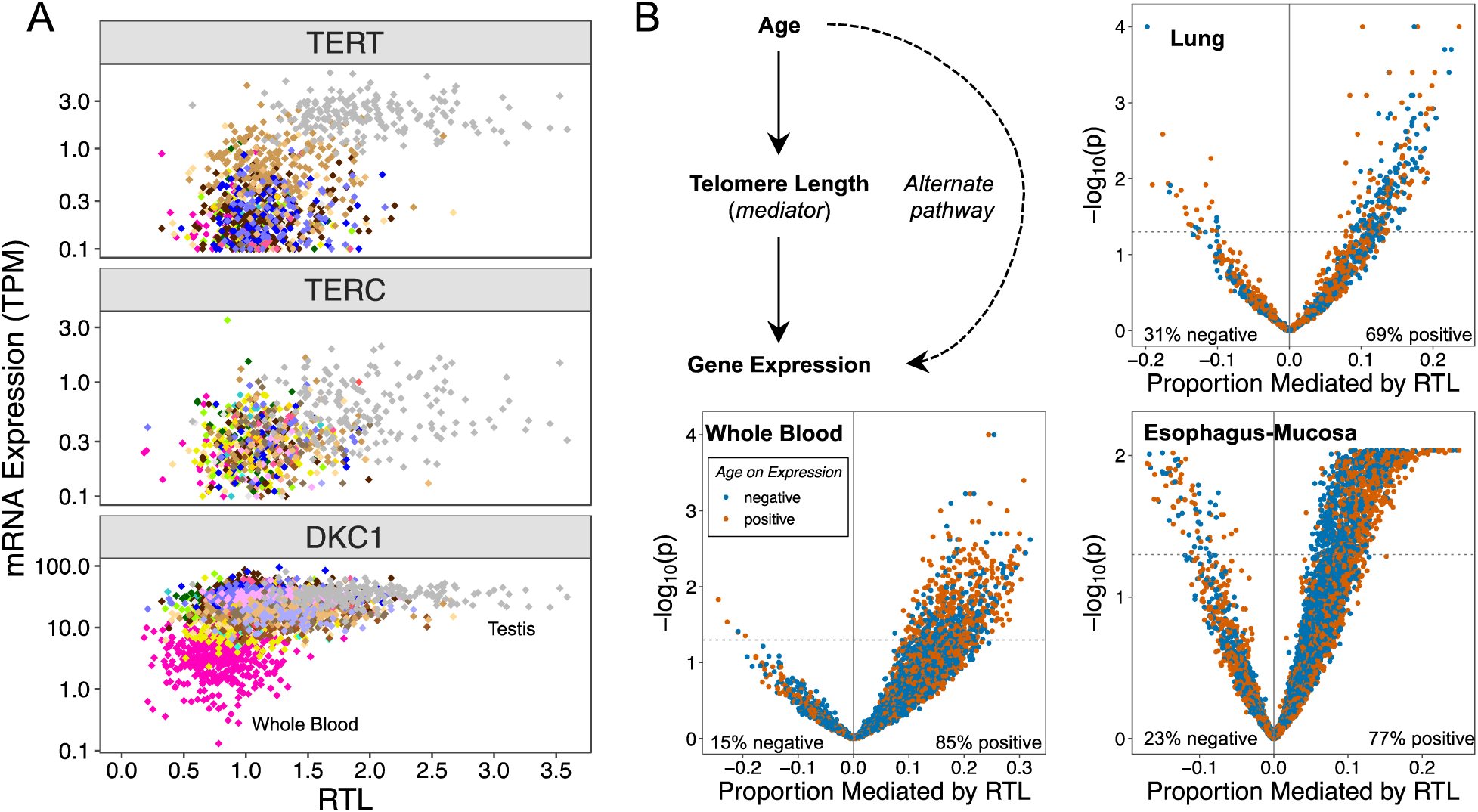
Telomere length is associated with telomerase subunit gene expression and mediates the effect of age on gene expression. A) RTL plotted against *TERC, TERT*, or *DKC1* expression across tissue types. Colors correspond to GTEx tissue types. B) Analyses addressing the hypothesis that TL mediates the effect of age on expression of specific genes. Scatterplots show estimates of the proportion of the effect of age on gene expression mediated by RTL (for each gene) and the −log_10_(*p*-value) corresponding to the average causal mediation effect of RTL (for each gene). Results are presented for all age-associated genes in each of the three selected tissue types (whole blood, lung, and esophagus-mucosa [EM]). The mediation *p*-value was obtained using a nonparametric bootstrapping approach (n=10,000 bootstraps).

### TL is associated with telomerase subunit expression across tissues

The telomerase enzyme can extend the telomere repeat sequence, typically in stem and/or progenitor cells, to compensate for TL shortening. The protein products of *TERT, TERC*, and *DKC1* comprise the telomerase catalytic subunit. We examined the association between RTL and expression of these genes using 3,885 GTEx tissue samples with both RTL and RNAseq (v8) gene expression data. *TERT* and *TERC* expression was detectable (i.e., transcripts per million [TPM] > 0.1) in 28% (n=1089) and 20% (n=783) of these samples (**Table S8** and **Table S9)**, respectively, but *DKC1* was ubiquitously expressed (n=3,885) in all samples (**Table S10**). While *DKC1* showed correlation with both *TERT* (r=0.30, *p* < 2 × 10^−16^, n=1089) and *TERC* (r=0.23, *p=*3 × 10^−11^, n=783) across all samples, the correlation between *TERT* and *TERC* expression across samples was stronger (r=0.49, p < 2 × 10^−16^, n=364) (**Figure S13**). Testis had substantially higher mean expression of *TERT* and *TERC* compared to all other tissues (p < 2 × 10^−16^) (**Table S8** and **Table S9**), but there was no association between testis RTL and *TERT* or *TERC* expression. Across all tissues, RTL was positively correlated with *TERT* (r=0.58, *p* < 2 × 10^−16^, n=1089), *TERC* (r=0.33, *p* < 2 × 10^−16^, n=783), and *DKC1* (r=0.29, *p* < 2 × 10^−16^, n=3,885) (**Figure 5A**). When testis was removed, the correlation decreased substantially for both *TERT* (r=0.14, *p=*4 × 10^−5^, n=890) and *DKC1* (r=0.23, *p* < 2 × 10^−16^, n=3,686) and disappeared for *TERC* (r=0.02, *p=*0.63, n=617). After adjustment for covariates and random effect of tissue type, RTL showed a positive association with increasing quartiles of *TERT* expression (*p=*0.005 including testis and *p=*0.002 excluding testis) and of *DKC1* expression (*p=*0.001 including testis and *p=*3 × 10^−4^ excluding testis) across all tissues. Overall these results support the following: (1) high telomerase activity in testis (i.e., spermatocytes) likely contributes to longer TL observed in that tissue and (2) GTEx tissue samples consist primarily of differentiated cells, which typically have little to no telomerase activity, resulting in minimal detectable association between telomerase activity in those cells and the observed TL (*32, 33*).

### TL mediates the effect of age on gene expression

Aging affects gene expression, and we sought to examine whether TL mediates the association between age and expression of age-associated genes. We analyzed the association between age and RNAseq-based gene expression levels among tissues with ≥ 150 samples, and selected three tissue types with >1,000 age-associated genes (FDR of 0.05) (see **Methods**): WB (n=5,153), lung (n=1,355), and EM (n=5,581) (**Figure 5B**). Using mediation analysis (*34*), we estimated the proportion of the effect of age on expression that was mediated by TL for each age-associated gene. For each tissue type, we observed substantially more positive than negative estimates of the “proportion mediated” (**Figure 5B**), as expected under the hypothesis that TL is a mediator (an equal number of positive and negative estimates are expected under the hypothesis of no mediation). We observed evidence that RTL mediated the effect of age on expression for 598 genes (12%) in WB, 224 genes (17%) in lung, and 1,108 (20%) in EM (based on p_mediation_ < 0.05 and proportion mediated > 0) (**Data Tables 4-6**). In these tissue types, RTL mediated between 4-32% of the effect of age on expression of individual genes; however, full mediation will be detected as partial mediation in the presence of measurement error (for either the mediator or the outcome) (*35*). We evaluated the enrichment of these RTL mediating genes in Gene Ontology (GO) terms (threshold p_enrichment_ < 10^−3^), and enriched GO terms were identified for lung (22 terms), WB (147 terms), and EM (104 terms) (**Data Tables 7-9**). Four terms were common to both WB and EM, and these were related to translation initiation and cellular adhesion. Six terms were common to lung and EM, and these were related to regulation of cell communication, regulation of signaling, calcium ion binding, cell periphery, and intrinsic and integral component of membrane. Among the 147 enriched GO-terms in WB, several terms related to apoptosis and cell death were identified.

### Tissue-level stem cell features are associated with TL and TERT expression

After extracting tissue-specific estimates of the number of division per stem cell (per year) and the proportion of stem cells (among all cells) for specific tissue types from Tomasetti and Vogelstein et al. (*36, 37*), we examined their relationship with mean RTL and mean *TERT* expression among non-reproductive GTEx tissue types (n=12, **Table S11**). No associations were identified between mean *TERC* and *DKC1* expression and these stem cell features. Mean RTL was positively correlated with estimated proportion of stem cells within a tissue type (r^2^=0.50, *p=*0.01) (**Figure 6** [top, left panel]), and this association persisted after adjustment for number of divisions per stem cell (*p=*0.02) and mean *TERT* expression (*p=*0.02). We did not observe a clear association between mean *TERT* expression and the estimated proportion of stem cells within a tissue type (**Figure 6** [top, right panel]). These results suggest that tissue types with a higher proportion of stem cells in their cellular composition may have longer TL measurements in bulk tissues as a consequence.

**Figure 6.**
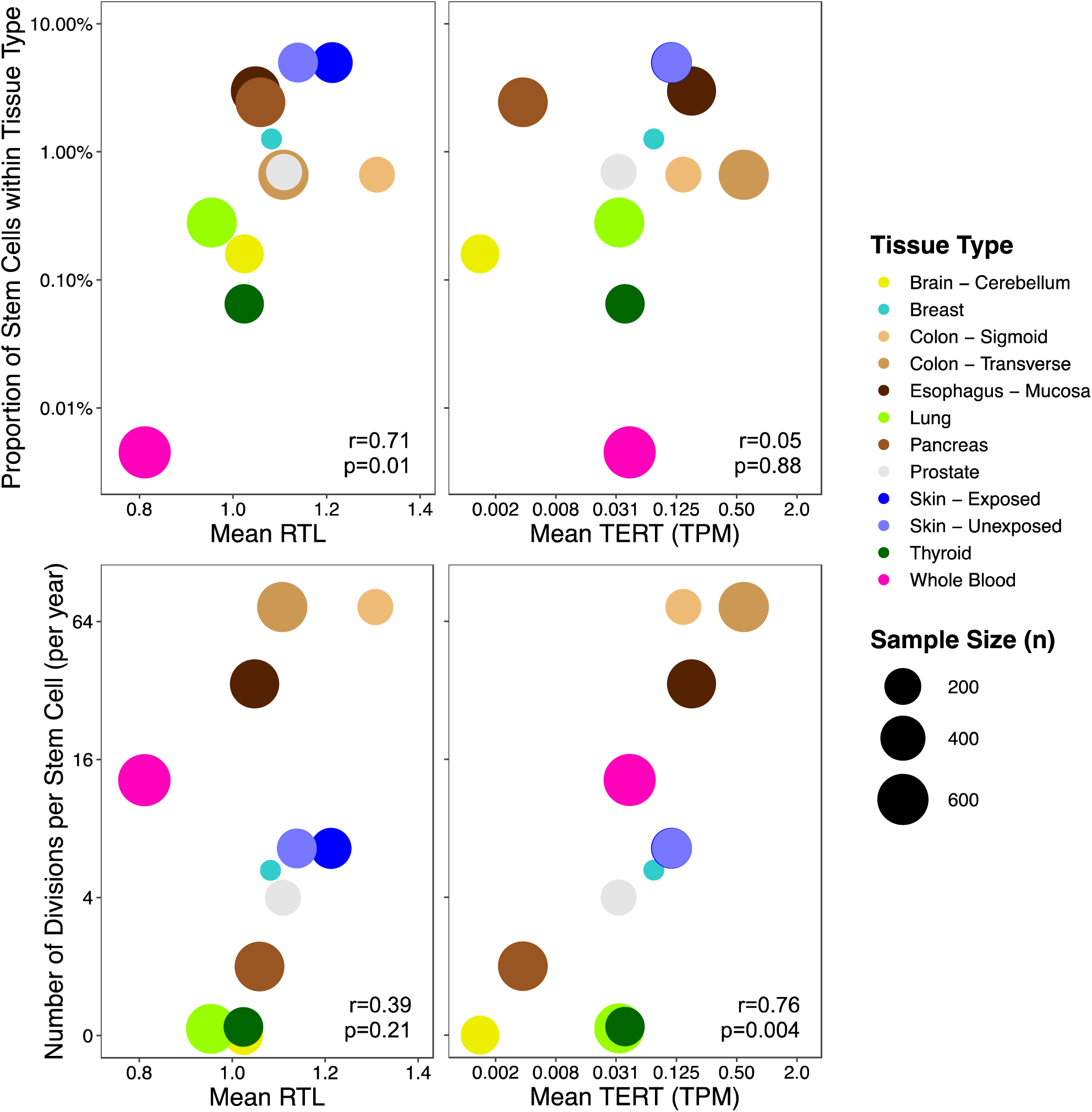
Telomere length and TERT expression are associated with estimated stem cell features. Estimated proportion of stem cells within tissues and its relationship between mean RTL (left) and mean *TERT* expression (right) are presented in top panel. Estimated number of divisions per stem cell (per year) within tissues and its relationship between mean RTL (left) and mean *TERT* expression (right) are presented in bottom panel. Colors correspond to GTEx tissue types, and size of points reflects sample size of tissue type. Pearson correlations and corresponding *p*-values are reported. Results are shown for non-reproductive tissues only.

We observed a positive correlation between mean *TERT* expression and the number of divisions per stem cell (r^2^=0.58, *p=*0.004) (**Figure 6** [bottom, right panel]). This association persisted after adjustment for the proportion of stem cells within a tissue type (*p=*0.007) and mean RTL (*p=*0.01). Mean RTL showed suggestive evidence of correlation with the number of divisions per stem cell (r^2^=0.15, *p=*0.21) (**Figure 6** [bottom, left panel]), and when we restricted to non-blood tissue types, mean RTL was positively correlated with number of divisions per stem cell (r^2^=0.42, *p=*0.03). This finding suggests that tissue types that undergo more cellular turnover and replacement, such as colon, may have higher telomerase expression in order to maintain TL in the stem cell compartments.

### Cell type composition is associated with TL within tissues

To determine whether TL varies among the cell types within a given tissue sample, we examined the association between RTL and estimated cell type enrichment scores (CTES) (generated using RNAseq data and the xCell software (*38*)). Seven CTES (for adipocytes, epithelial cells, hepatocytes, keratinocytes, myocytes, neurons, and neutrophils) were benchmarked by the GTEx consortium (Kim-Hellmuth et al 2019 [GTEx cell type paper]), and we examined the association between these 7 CTES and RTL in tissue types with ≥ 100 samples (n=16 tissue types). After removing cell types that were not detected within a tissue type (n=37 total CTES across 16 tissue types) and adjusting for age and sex, we identified eight associations (*p* < 0.05) between CTES and RTL among 37 associations tested (**Figure S14**). In exploratory analyses, we examined all 64 CTES provided by xCell that had a detection *p*-value < 0.05 for >90% samples within a tissue type. Restricting to tissue types with ≥ 300 samples that had both CTES and RTL data (WB, lung, and EM), there were 27, 24, and 17 CTES detected in each tissue, respectively (**Figure S15**). EM and lung had 13 and 14 CTES that were associated with RTL, after adjustment for age and sex (*p* < 0.05). RTL was positively associated with epithelial cell, smooth muscle cell, keratinocyte, and sebocytes CTES in both lung and EM (*p* < 0.05). Notably, five CTES were inversely associated with RTL (*p* < 0.05) in both lung and EM, including fibroblasts and endothelial cells. In WB, lymphoid and myeloid cell CTESs accounted for 70% of the CTES detected, and eight CTES were associated with RTL, respectively (*p* < 0.05). Neutrophil CTES were positively associated with RTL. Both CD8+ T-cell CTES were inversely associated with RTL, consistent with prior work examining cell types and TL in blood (*39*). These results provide evidence that TL varies across cell types within a given tissue, and consequently, cell type composition can affect TL measurement in human tissues.

### TL across all tissues is associated with age-related chronic disease status

Using medical history data from GTEx donors, we examined the association between common age-related chronic diseases and RTL within and across tissues. A history of Type II diabetes (22% of donors) was associated with shorter RTL across all tissues (*p=*0.02), as well as, shorter pancreas RTL (*p=*0.07) and coronary artery RTL (*p=*0.01) (**Figure S16**). Among all donors, 50% had no history of any chronic disease, and 30%, 14%, and 6% had a history of one, two, and three (or more) chronic diseases, respectively. Chronic disease burden (sum of chronic diseases from 0-5) was associated with shorter RTL across all tissues (*p=*0.008) and in testis (*p=*0.03), coronary artery (*p=*0.03), kidney cortex (*p=*0.04), and cerebellum (*p=*0.009). When we excluded cancer from the chronic disease burden, these associations persisted across all tissues (*p=*0.02) and in all tissues listed above except for kidney cortex (*p=*0.09). These observations suggest that TL may capture some aspect of the biologic age-related health decline across tissues.

We did not observe any associations between RTL and history of cancer; however, to test the hypothesis that normal tissues with relatively short (or long) TL are also short (or long) in tumors occurring in that tissue, we compared the mean tissue-to-WB TL ratio for each GTEx tissue to the mean tumor-to-WB TL ratio in corresponding cancer types from The Cancer Genome Atlas (TCGA) (see **Methods**) (*40*). Mean cancer TL ratio from TCGA and normal TL ratio from GTEx were positively correlated (r=0.44, *p=*0.03, n=23) (**Figure S17**), providing support for this hypothesis.

## Discussion

This study provides an unprecedented view of the substantial variation in human TL that exists across human tissue types and among individuals. We show that TL is generally positively correlated across human tissue types, and that whole blood TL can serve as a proxy for tissue-specific TL for many tissues, a finding that may support the use of blood TL as a proxy for TL in some tissues in large epidemiological studies. TL was negatively associated with age in the majority of tissues studied, confirming the hypothesis of pervasive age-related telomere shortening in most human tissues. However, the rate of shortening varied across tissues, and age explained more variation in TL in tissues with shorter mean TL. *TERT* and *TERC* expression were low or undetectable in most tissues and were not associated with TL within any tissue, likely because progenitor cells, which express telomerase, are not present in large numbers in adult tissues, which consist primarily of differentiated cells. Notably, testicular TL was ∼1.5-2.5-fold longer than TL in any other tissue type, and *TERT* was expressed in 100% of these samples and at higher levels than any other tissue, consistent with predominance of spermatogenic cells in testis (i.e., cells developing from germ cells into spermatozoa) which have high telomerase activity (*33*).

RTL measured in a tissue sample is an average of the TL among all chromosomes within a heterogeneous population of cell types with different cell division rates/history, stem cell composition, and oxidative and inflammatory environments. In order to characterize variation in TL within specific cell types, cell type-specific and single-cell TL studies are needed. A large proportion of the variation in RTL was unexplained across all tissue types, potentially attributed to sources such as cell type composition (e.g., stem and progenitor cells), measurement error, and lifestyle and environmental factors with variable effects across tissues. From our simulation-based analysis of the impact of TL measurement error on our results, we show that random measurement error biases our estimate of the true correlation in TL between two tissues towards zero, suggesting that the correlations presented in this study are attenuated compared to the true correlations. We lack detailed monitoring to exposure data (e.g., smoking and alcohol use) for GTEx donors; studies that can link human tissue samples to environmental and lifestyle histories are needed to better understand environmental determinants of TL across different tissues and cell types. Currently, all TL-associated SNPs have been identified in genome-wide association studies of leukocyte TL (*12-15*); our study suggests some of these effects are also present in other tissue types, but larger studies of tissue-specific TL measurements are needed to characterize how these effects vary across tissues and cell types. Identifying variants that impact TL in all or most cell types (e.g., variants with effects on TL that may be present during development or in stem cells in multiple tissue types) may be ideal for evaluating the causal impact of TL on risk for a wide array of diseases (occurring in diverse tissues or cell types) using Mendelian randomization. Future studies should also evaluate the relationship between TL and other biologic and cellular aging processes and biomarkers of aging within and across tissues in order to further characterize the role TL in human aging.

## ACKNOWLEDGEMENTS

We would like to acknowledge the GTEx donors and families for their generous participation in and contribution to the GTEx consortium. We thank A. Aviv for his helpful comment and review of the manuscript.

## Author contributions

K.D., J.D., L.S.C., M.G.K, H.A., B.L.P. conceived and designed the study. F.J., J.S., M.S., M.G.K. conducted Luminex assays on all samples. K.D., L.S.C., M.C., L.T., H.L., E.R., B.L.P. contributed to the statistical analyses in the study. K.A. and F.A. were responsible for the generation of GTEx v8 RNAseq and genotyping data for the GTEx consortium. M.O., S.K.H., B.E.S., generated the GTEx cell type estimates for the v8 release. K.D. and B.L.P. wrote the manuscript. All authors contributed to the revision and review of the manuscript.

## Data and materials availability

All data will be available on the GTEx Portal (www.gtexportal.org) and deposited on dbGap for future research use.

