## Supplementary material for "Determinants of telomere length across human tissues": Methods

#### MATERIALS AND METHODS

***GTEx tissue donors and tissue samples***. A detailed description of GTEx recruitment has been published previously (*20, 41*). Briefly, all GTEx study participants were post-mortem tissue donors. Donors were excluded if they had a BMI >35 or BMI <18.5, metastatic cancer, or potential exposure to human immunodeficiency virus (HIV) or other viral infections. Demographic, death report, and medical history questionnaires were completed for all GTEx donors by the Comprehensive Data Resource, using information from medical records, next of kin, and other sources. Most tissue samples were obtained from each donor within a < 8 hour postmortem interval (PMI) (*41*). All tissue samples were preserved using the PAXgene tissue preservation system (Qiagen). Venous whole blood was obtained and preserved in ACD (acid citrate dextrose) and PAXgene blood tubes (Qiagen). DNA and RNA were isolated and stored at -20C and -80C, respectively (*20*). Consent to participate in the study was obtained from immediate family members and details are described by Siminoff et al (*42*). The research protocol was reviewed by Chesapeake Bay Review Inc, Roswell Park Cancer Institute's Office of Research Subject Protection, and the Institutional Review Boards (IRB) at the University of Pennsylvania. Analyses of GTEx DNA samples at the University of Chicago was not considered human subjects research by the IRB at the University of Chicago because only de-identified data on deceased individuals were involved.

***Relative Telomere Length (RTL) Measurement***. GTEx tissue samples were stored at the GTEx Laboratory and Data Analysis Coordinating Center (LDACC) at the Broad Institute, and DNA was extracted using the Qiagen Gentra Puregene method for GTEx tissue samples included in this study. DNA samples were sent to the Institute for Population and Precision Health Laboratory at the University of Chicago (directed by Dr. Kibriya) on 96-well plates, with 64 samples on each plate. Each plate contained at least one replicate sample. We measured RTL for all samples using a high-throughput, probe-based assay for RTL measurement that uses QuatiGene chemistry on a Luminex platform (*22, 43-45*). Among the GTEx samples measured in duplicate for RTL, the assay precision was good to excellent, with an intraclass correlation coefficient (ICC) of 0.91 (95% CI 0.86–0.94). Among all GTEx samples measured, the correlation with age was -0.41, providing external validation for the assay. We used a modified version (*22, 45*) of the original Luminex assay (*44*) to increase the throughput. Briefly, the assay requires ~50 ng of DNA, which is hybridized to sequence-specific probes for the telomere repeat sequence (TEL) and reference gene sequence (ALK). The TEL and ALK gene signals are amplified using branched DNA technology and detected using Luminex technology. RTL is expressed as the telomere quantity index (TQI), calculated as TEL/ALK. Therefore, TQI is a measure of *relative* TL (RTL), which is normalized for the quantity of DNA in a well, and it is relative to a "standard" DNA sample (*45*). This Luminex assay requires a small quantity of input DNA, involves no DNA amplification, does not show well position effects (*45*), and can efficiently measure RTL in many samples. The Luminex assay has been validated against the Southern blot method in a blinded comparison, showing a correlation with Southern blot TL in the range of 0.65-0.75 (*22*).

Samples were assigned to a plate based on their DNA concentration (high, medium, or low), which prevented complete randomization of tissue types across plates but tissues were not batched by plate. All plates consisted entirely of samples from one of the three concentration groups in order to enable consistent re-concentration of samples for RTL measurement. Among the 126 plates (7,535 samples) for which we attempted measurement, five low concentration plates with 63 samples each failed and 301 samples were removed from the analysis. Among the remaining plates and samples (n=7,234), RTL measurement failed for 836 samples (11.6%), due to very low intensities and/or low bead count mainly from DNA samples with low concentration.

We analyzed 243 replicate samples and observed a strong positive correlation between the two RTL measurements (Pearson's r=0.85, $p < 2.2 \times 10^{-16}$) (**Figure S18**). We estimated TL from available WGS data for whole blood samples in GTEx (n=443) using the Telseq software (*46*) in order to compare our RTL measurements in GTEx. The correlation (r) between these two TL measures was 0.30 (**Figure S19**). Among all tissue types, technical factors, represented by plate (e.g. batch effects, DNA quality and concentration), and donor ischemic time, explained 11.2% and 3.7% of the variation in RTL, respectively. We adjusted for these factors in the analyses described below.

***Processing and quality control for RTL data***. We examined the distribution of RTL within each tissue type and detected potential outliers by calculating the interquartile range (IQR) for each tissue type and flagging samples that were greater or less than 3*IQRs from the median. We identified seven outliers (≥ 3*IQRs from median) and removed these samples within the following tissues: thyroid, prostate, brain (cerebellum), esophagus (gastric junction), and skin (unexposed). After these exclusions, we then examined the distribution of RTL for each tissue and computed the mean and standard deviation (SD).

***Within and across tissue analyses and estimation of variance explained in RTL***. Age, sex, and race/ethnicity were tested for association with RTL for tissue types with ≥ 75 samples. Pearson correlations and *t*-tests were used to examine unadjusted relationships between RTL and covariates. Within each tissue type, we ran linear mixed models (LMMs) adjusted for age, sex, race/ethnicity, BMI (continuous), donor ischemic time, and technical factors [DNA concentration and sample plate] as a random effect. We used the lme4 package in R to perform LMM and obtained *p*-values from the LMMs using likelihood ratio tests (LRT) comparing the model with and without the covariate of interest.

***Estimation of variance explained in RTL.*** For analyses across all tissue types, we used LMMs to estimate the contribution of covariates to variance explained in RTL. The PVE (percent variance explained) for the combined fixed-effect covariates (age, sex, donor ischemic time, BMI, race/ethnicity, and technical factors) and random effects (tissue and donor) were estimated by extracting the variance components for the combined fixed effects and each random effect and dividing by the sum of all variance components (fixed effects, random effect for tissue type, random effect for donor, and residual). To estimate the contribution of each fixed effect to the variation explained in RTL, we used the proposed approaches by Nakagawa et al. and the *MuMIn* package in R (*47*). For each fixed effect of interest, we ran an LMM that included a single fixed effect and the random effects of tissue type and donor. We estimated the contribution of variance explained (%) by that fixed effect across all tissue types and donors from the marginal $R^2$ (which captures variance explained by fixed effects) and extracted the *p*-value from a LRT comparing the model with the fixed effect to the model without the fixed effect.

***Relationship among RTL measures from different tissue types***. We computed Pearson correlations between tissue type pairs among tissue types with available RTL data for ≥ 75 samples. In order to cluster tissues by strength of RTL co-variability, hierarchical clustering analysis was applied using the pairwise correlations as our distance measure and average linkage to determine clusters (*48*). Between whole blood RTL and tissue RTL pairs, we also computed partial correlation adjusted for age, sex, race/ethnicity, BMI, and donor ischemic time.

In order to examine the influence of non-differential measurement error in RTL measurements, we generated simulated data to examine how random error (i.e., noise) in RTL affects the observed Pearson's *r* between RTLs from two tissue types. For each simulated dataset, we randomly generated zygote RTLs from a standard normal (0,1) distribution (under the assumption that zygote TL is the primary diver of correlation in TL among adult tissues). Tissue-specific RTLs were generated according to the following equation: $RTL = (\beta \times zygote\ RTL) + (\sqrt{1-\beta^2} \times \varepsilon)$ with $\varepsilon \sim N(0,1)$. The β value was varied in order to yield specific *r* values (0.2, 0.4, 0.6, and 0.8) representing the true correlation between the TL values of the two tissues. These "error-free" TL variables had standard normal (0,1) distributions in each tissue. Measurement error for each tissue was randomly-generated based on a normal (0, $sd_{me}$) distribution, where the standard deviation of the measurement error component ($sd_{me}$) was varied in order to generate a specific proportion of variance attributable to measurement error ($p_{me}$) using the following formula: $sd_{me} = \sqrt{p_{me}/(1-p_{me})}$. The $p_{me}$ values evaluated were 0%, 10%, 25%, and 50%. This error was added to the simulated "true" tissue TL values to generate the "observed" TL values. For each scenario (defined by a true *r* value and the proportion of variance due to measurement error in each tissue) we generated 1000 datasets, each with 500 observations. We estimated the r between the tissue-specific RTL variables for each simulated dataset and extracted the median *r* for each scenario (see **Figure S6**).

We also analyzed within-person differences in RTL between tissue types (ΔRTL) and the correlation of these differences with age. We restricted our analysis to pairs of tissues with a sample size ≥ 50. Among the 155 pairs analyzed, we examined the Pearson correlation between ΔRTL and age and reported the absolute value of the correlation (since correlation depended on order of tissues when computing the difference). Linear models also were examined the relationship between age and ΔRTL adjusted for sex, race/ethnicity, BMI, and donor ischemic time.

***Association between RTL and a polygenic score for leukocyte TL***. We identified genetic variants (SNPs) associated with leukocyte TL from prior genome-wide association studies (*12-14).* We selected nine SNPs that showed consistent associations with leukocyte TL across studies and were not in linkage disequilibrium with each other: rs11125529, rs10936599, rs2736100, rs7675998, rs9420907, rs2535913, rs3027234, rs8105767, and rs755017 (**Table S8**). We extracted the association estimates for these SNPs from the ENGAGE consortium GWAS (*12*). The polygenic SNP score for TL was estimated as a weighted linear combination of TL-increasing alleles, corresponding to the sum of the weighted allele counts across all 9 SNPs ($\sum_{i}^{n=9} \beta_{LTL\_sd} \times allele\ count$). We included tissues with RTL and genetic information for ≥ 100 samples. Within tissues, the relationship between SNP score and RTL was examined using linear models adjusted for age, sex, donor ischemic time, the first five genotyping PCs, and a random effect representing technical factors (DNA concentration and sample plate). LMMs were used to examine the relationship between SNP score and RTL across all tissues types and adjusted for fixed covariates from linear models and random effect of tissue and donor.

***Co-localization analyses for leukocyte TL-associated loci and eQTLs***. We used co-localization analyses to assess the evidence that TL-associated loci shared a common causal variant with eQTLs observed based on GTEx eQTL (v8) (GTEx Consortium 2019 [GTEx main paper]) using the Bayesian test for co-localization proposed by Giambartolomei et al. and implemented in the *coloc* software (*49*). We extracted the summary statistics for the nine leukocyte TL-associated regions from the ENGAGE consortium (*12*) and remapped SNPs to hg38. To identify potentially co-localizing eQTLs, we first determined the tissues for which the nine TL-associated SNPs were listed as cis-eQTL SNP based on GTEx v8 results. From these results, we selected tissue-specific eQTLs for co-localization analysis if the lead TL-associated SNP was associated with expression with a $-\log_{10}$(p-value) that was > 0.9 when divided by the $-\log_{10}$(p-value) for the lead SNP of the cis-eQTL. For cis-eQTLs meeting this criteria, we conducted co-localization analyses using the summary statistics from the ENGAGE GWAS of leukocyte TL and GTEx v8 cis-eQTL results.

Co-localization analysis requires specifying prior probabilities for a SNP being causal for expression only (p1), TL only (p2), and both traits (p12). We first made the assumption that the overall probability of a SNP a being causal for gene expression (eQTL) was $10^{-4}$ and the probability of a SNP being causal for TL was $10^{-5}$. With these assumptions in place, we then created three sets of co-localization parameters corresponding to 50%, 20%, and 5% of causal TL SNPs affecting gene expression. These corresponding prior sets for the co-localization parameters were: p1=$9.5\text{x}10^{-5}$, p2=$5.0\text{x}10^{-6}$, p12=$5.0\text{x}10^{-6}$ (50%); p1=$9.8\text{x}10^{-5}$, p2=$8.0\text{x}10^{-6}$, p12=$2\text{x}10^{-6}$ (20%); and p1=$9.9\text{x}10^{-5}$, p2=$9.5\text{x}10^{-6}$, p12=$5\text{x}10^{-7}$ (5%). The *coloc* software provides a posterior probability of a shared common causal variant for each co-localization test conducted (PP of CCV or PP (H4) in **Data Table S3**).

***Association of RTL with expression of TERT, TERC, and DKC1***. The GTEx project has generated RNAseq data for the vast majority of GTEx tissue samples, as described in Mele et al. (*50*). Briefly, RNA samples with a RIN >=6 were sequenced, expression was quantified using RNA-SeQC v1.1.8, and transcript level quantifications were calculated using RSEM v1.2.22 (*51*). Gene expression data was obtained for GTEx donors included in the GTEx v8 release (n=3,885 with both RNAseq and RTL data).

We extracted TPM (transcripts per million) expression measurements for *TERT* (ENSG00000164362), *TERC* (ENSG00000270141), and *DKC1* (ENSG00000130826) and used a threshold of TPM > 0.1 for detectable expression in our analyses. Pearson correlations were used to assess the unadjusted association of gene expression with age and RTL. Within tissues, we ran LMMs adjusted for age, sex, race/ethnicity, BMI (continuous), donor ischemic time, and technical factors [DNA concentration and sample plate] as a random effect. Across tissues, we modeled gene expression as quartiles (defined among samples with detectable expression) and examined the relationship between RTL and gene expression in a LMM adjusted for fixed effect covariates and tissue type as a random effect.

***Mediation analyses of age-associated gene expression***. Using the RNAseq tissue-specific TMM normalized and standardized gene expression datasets (all genes expressed on standard normal (0,1)) and corresponding covariate files (GTEx v8), we identified age-associated genes within specific tissue types using linear models adjusted for age, sex, donor ischemic time, RNAseq platform, and first five genotyping PCs. We extracted expression data for age-associated genes for three tissues that had ≥ 100 age-associated genes (FDR of 0.05): whole blood, lung, and EM. For each age-associated gene identified, we conducted mediation analysis using the *mediate* R package (*34*), to assess the evidence that the effect of age on gene expression is mediated by TL. We adjusted for the same covariates in the mediator and outcome models (age, sex, genotyping PCs [1-5], RNAseq platform, and donor ischemic time) and ran 10,000 bootstraps using the non-parametric method to obtain p-values and 95% confidence intervals for the average causal mediation effect (i.e., indirect effect), direct effect, and total effect. The average causal mediation effect provides an estimate of indirect effect of age on age-associated gene expression that is mediated by TL. The "proportion mediated" was obtained by dividing the average causal mediation effect by the total effect, reflecting the extent to which adjustment for TL attenuates the association between age and expression of a given gene. Among the genes showing evidence of mediation by TL ($p < 0.05$) for each tissue type, we conducted geneset enrichment analysis (GSEA) among Gene Ontology (GO) terms using the *goana* function in *limma* and extracted all terms with $p < 0.001$ (from Fisher's exact test), restricting to GO terms with 3 or more TL-mediating genes and between 10-5000 genes in pathway.

***Tissue-specific stem cell features***. We obtained tissue-specific estimates of the number of stem cell divisions and the total normal and stem cell population from prior Tomasetti and Vogelstein publications (*36, 37*). We summed the stem cells for the following tissues: pancreas (ductal and endocrine cells), thyroid (follicular and parafollicular), and skin (basal and melanocytes) and constructed weighted average for stem cell division rate for skin (40% melanocytes and 60% basal). The stem cell division rate for cell types in pancreas and thyroid were the same. We computed the proportion of stem cells within a tissue from the total stem cells divided by the total normal and stem cells. These stem cell estimates were available for 14 GTEx tissue types, and we restricted our analysis to non-reproductive tissue types (n=12), excluding testis and ovary (**Table S11**). We examined Pearson and Spearman correlations between stem cell features (i.e., number of divisions per stem cell and proportion of stem cells) of a given tissue and the mean RTL and mean *TERT* expression of those tissues (n=12).

***Generation of cell type enrichment scores***. The xCell software was used to generate cell type enrichment scores (CTES) for each GTEx tissue sample (Kim-Hellmuth et al 2019 [GTEx cell type paper]), xCell is a gene expression signature-based method that estimates enrichments for 64 immune and stromal cell types (*38*). Cell type enrichment scores (CTES) were generated using RNA gene expression (expressed in TPM) data as input and single sample enrichment scores and detection p-values were determined for 64 cell types. Cell types with a CTES detection p-value > 0.05 (i.e., not present) in a sample were set to missing. Within each tissue type, we applied an inverse normal transformation to the CTES for each cell type. We first analyzed the associations between benchmarked CTES selected by the GTEx consortium (adipocytes, epithelial cells, hepatocytes, keratinocytes, myocytes, neurons, and neutrophils) and RTL adjusting for age and sex. We retained cell types that were detected in 75% or more samples within a tissue type (n ≥ 100) and had a median CTES > 0.01 within that tissue type. In an exploratory analysis, we included all 64 CTES in tissue types with ≥ 300 samples that had both CTES and RTL data (i.e., WB, lung, and EM). For each tissue type, we retained cell types that were detected (CTES detection p-value < 0.05) in 90% or more samples and had a median CTES > 0.01 within that tissue type. We assessed the association between RTL and CTES adjusting for age and sex.

***Association analyses of chronic disease history and RTL***. The GTEx project provides coded medical history data for all donors. We determined the top five causes of death from chronic disease among adults 20-70 years using the web-based CDC Wonder (http://wonder.cdc.gov). Based on ICD-10 coding, the top five causes of death from chronic disease from 1999-2016 were malignant cancer (30.4%), ischemic heart disease and other heart disease (19.0%), chronic lower respiratory disease (4.9%), cerebrovascular disease (3.7%), and type II diabetes (3.5%). We extracted the medical history variables for these five chronic diseases and recoded into binary variables. We created a chronic disease index (CDI) by summing these five variables (range 0-5), and collapsed individuals with values greater than two. As a sensitivity analysis, we also examined the index without cancer. Only seven individuals had a chronic disease index of 4 or 5. We examined the relationship between history of chronic disease and RTL using the within tissue LMM modeling and across all tissue types LMM, described above, and assessed both unadjusted and adjusted results.

***Comparison of TL estimates in GTEx tissue types to cancer types from The Cancer Genome Atlas (TCGA).*** We downloaded the available TCGA TL data as processed by Bartel et al (*40*). Briefly, the TL for cancer samples were estimated using the Tel-seq software (*46*) using whole genome sequencing, low pass sequencing, and exome sequencing data. We utilized the cancer TL to normal whole blood TL ratios, which reduced the influence of technical effect from sequencing center and method (*40*). Among our GTEx samples with RTL, we constructed normal tissue type TL to normal whole blood TL ratios. We extracted the mean ratios from TCGA from cancer types that corresponded to tissues included in the GTEx analysis (n=18 cancer types out of 31) and computed the mean ratio from GTEx for these corresponding tissues (n=18). We examined this relationship using Pearson's correlation.

All analyses were performed in R 3.5.2.

### Supplementary Materials for

#### Determinants of telomere length across human tissues

Kathryn Demanelis, Farzana Jasmine, Lin S. Chen, Meytal Chernoff, Lin Tong, Justin Shinkle, Mekala Sabarinathan, Hannah Lin, Eduardo Ramirez, Meritxell Oliva, Sarah Kim-Hellmuth, Barbara E. Stranger, Kristin G.Ardlie, François Aguet, Habibul Ahsan, GTEx Consortium, Jennifer Doherty, Muhammad G. Kibriya, and Brandon L. Pierce

**This PDF file includes:**

Figs. S1 to S19
Tables S1 to S11
Captions for Data S1 to S9

**Other Supplementary Materials for this manuscript include the following:**
Data S1. Pairwise Tissue RTL Pearson Correlations.

Data S2. Difference in RTL between tissue pairs (ΔRTL) and the association with age.

Data S3. Posterior probabilities of co-localization for telomere length association signals and *cis*-eQTLs.

Data S4. Mediation analyses assessing telomere length as potential a mediator of the effect of age on gene expression in whole blood.

Data S5. Mediation analyses assessing telomere length as potential a mediator of the effect of age on gene expression in lung.

Data S6. Mediation analyses assessing telomere length as potential a mediator of the effect of age on gene expression in esophagus - mucosa.

Data S7. Gene-set enrichment results for age-associated genes mediated by telomere length in whole blood.

Data S8. Gene-set enrichment results for age-associated genes mediated by telomere length in lung.

Data S9. Gene-set enrichment results for age-associated genes mediated by telomere length in esophagus - mucosa.

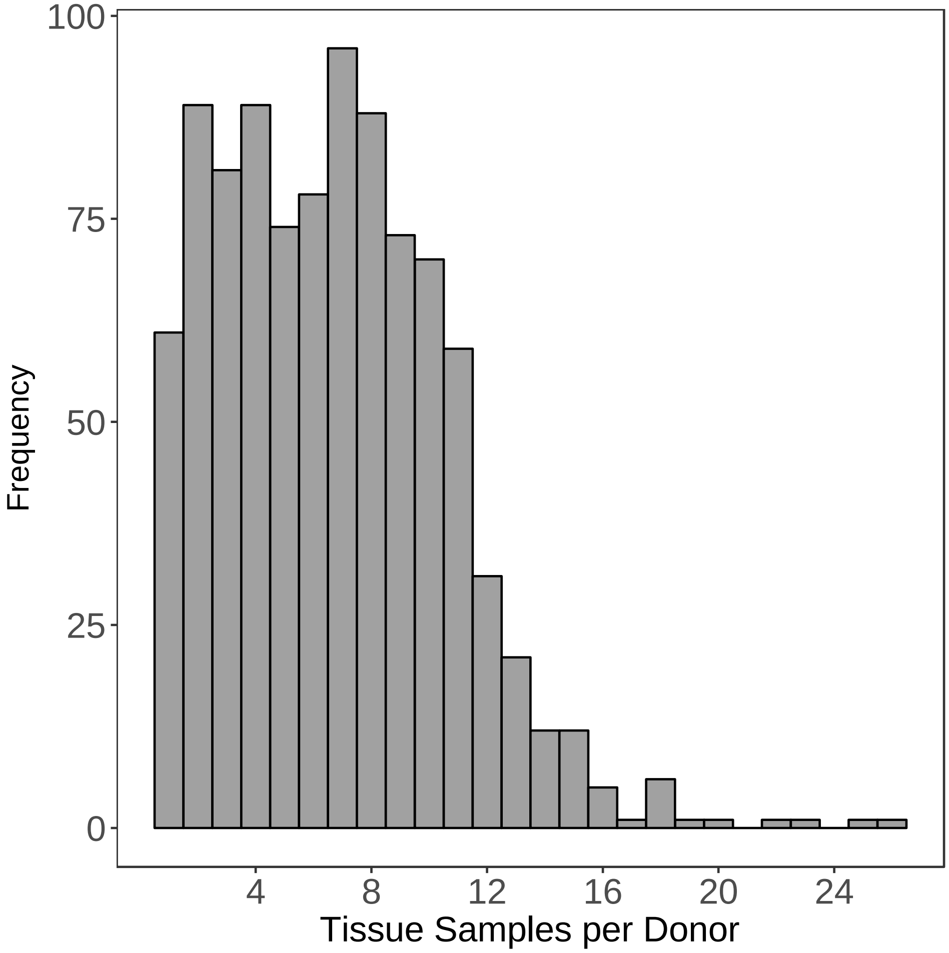


**Fig. S1.**
Distribution of the number of tissue samples per donor with data on telomere length.

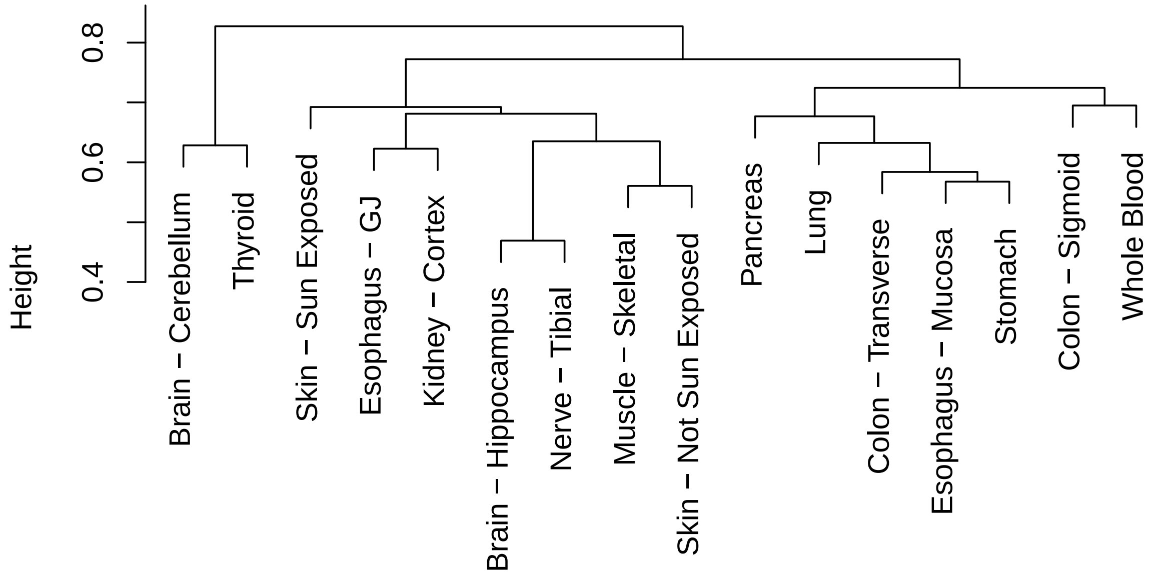


**Fig. S2.**

Dendrogram from hierarchical clustering analysis of tissue RTL measures from 16 tissues). Only tissues with ≥ 75 samples were included, and hierarchical clustering utilized average linkage and tissue-pair Pearson correlations for distance matrix. Corresponds to **Figure 2B**.

A.

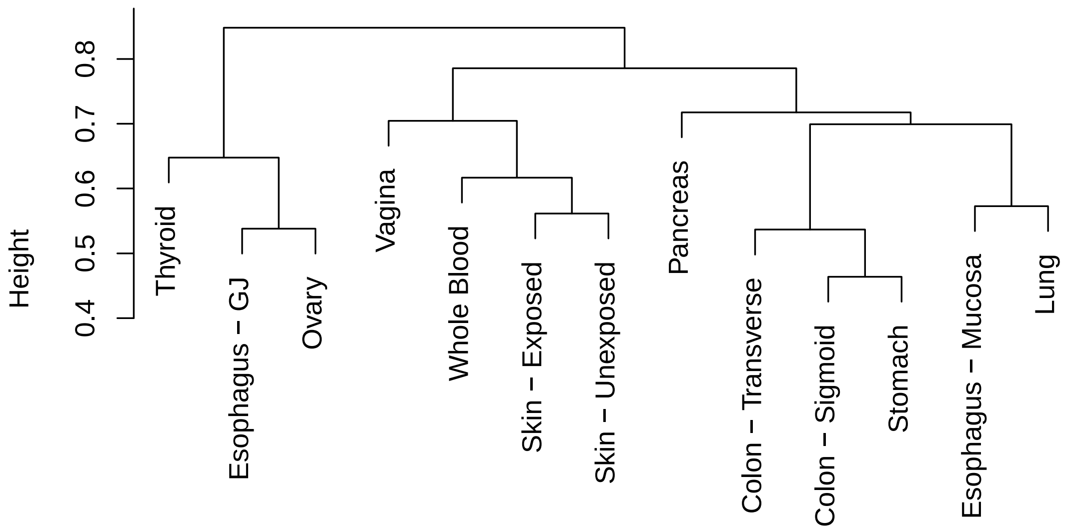


B.

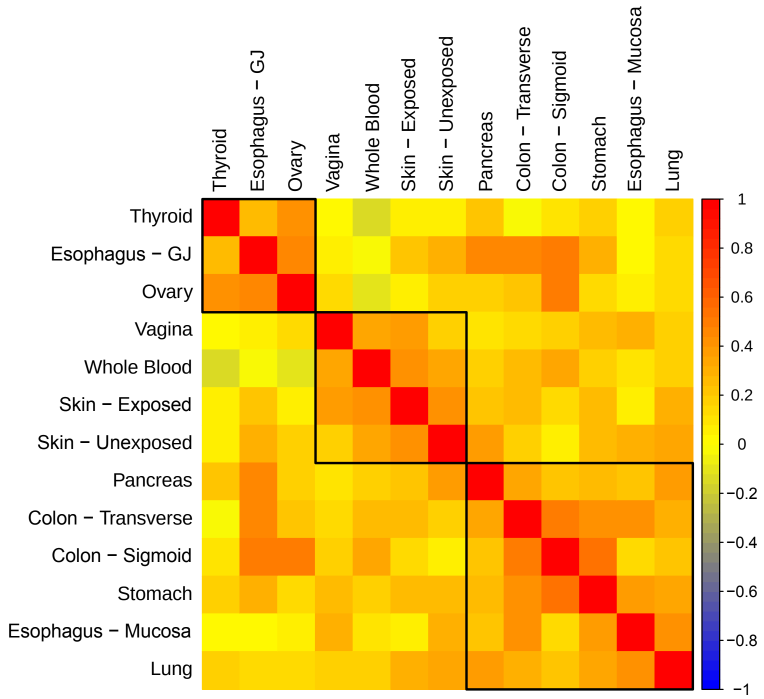


**Fig. S3.**

Clustering of RTL among tissues from females (n=13 tissues). Only tissues with ≥ 75 samples were included, and hierarchical clustering utilized average linkage and tissue-pair Pearson correlations for distance matrix. A) Dendrogram of tissue RTL from hierarchical clustering analysis among tissues from females. B) Tissue pair Pearson (r) correlations among tissues from females. Red, yellow, and blue correspond r=1, r=0, and r=-1, respectively. Black boxes are results from hierarchical clustering for k=3 clusters.

A.

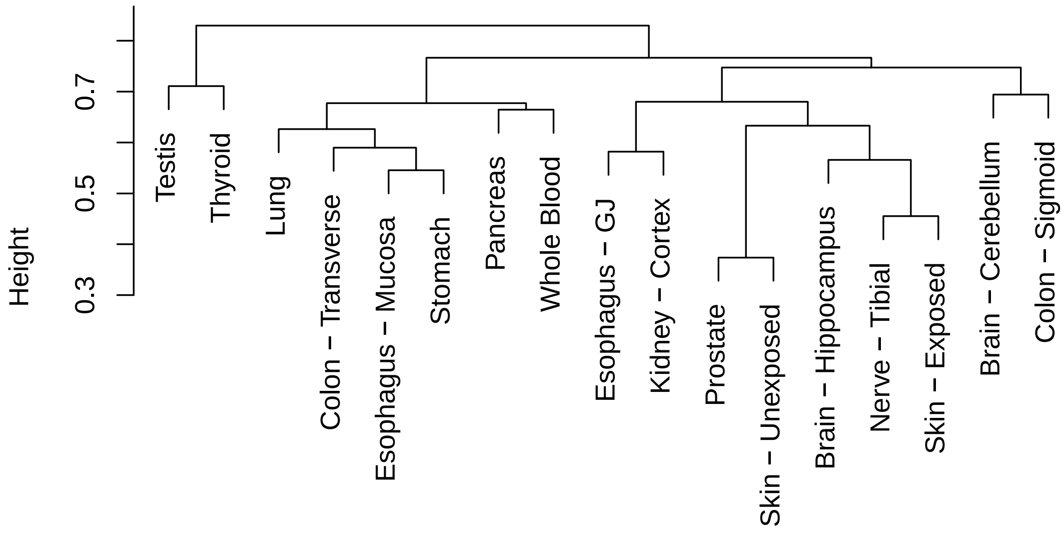


B.

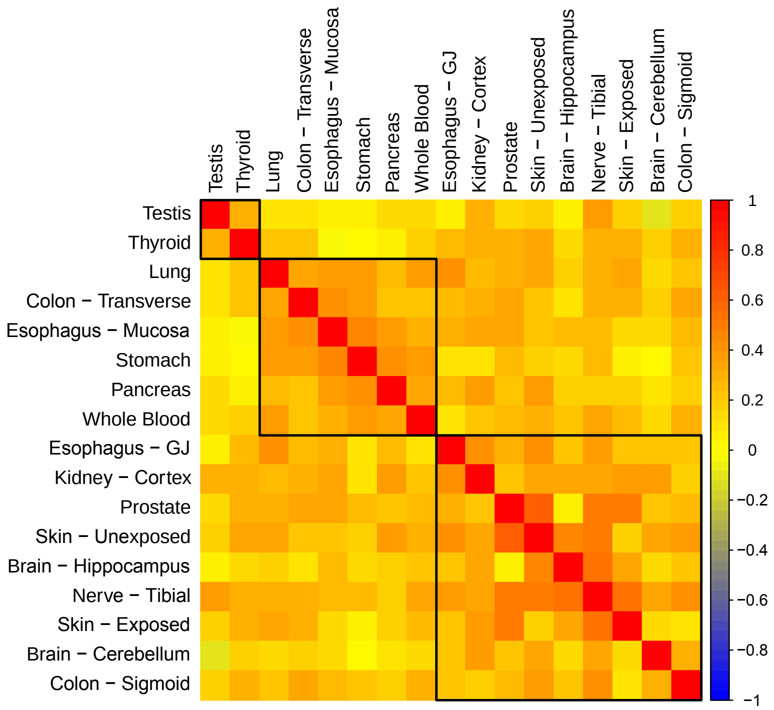


**Fig. S4.**

Clustering of RTL among tissues from males (n=17 tissues). Only tissues with ≥ 75 samples prior to stratification by sex were included, and hierarchical clustering utilized average linkage and tissue-pair Pearson correlations for distance matrix. A) Dendrogram of tissue RTL from hierarchical clustering analysis among tissues from males. B) Tissue pair Pearson (r) correlations among tissues from males. Red, yellow, and blue correspond r=1, r=0, and r=-1, respectively. Black boxes are results from hierarchical clustering for k=3 clusters.

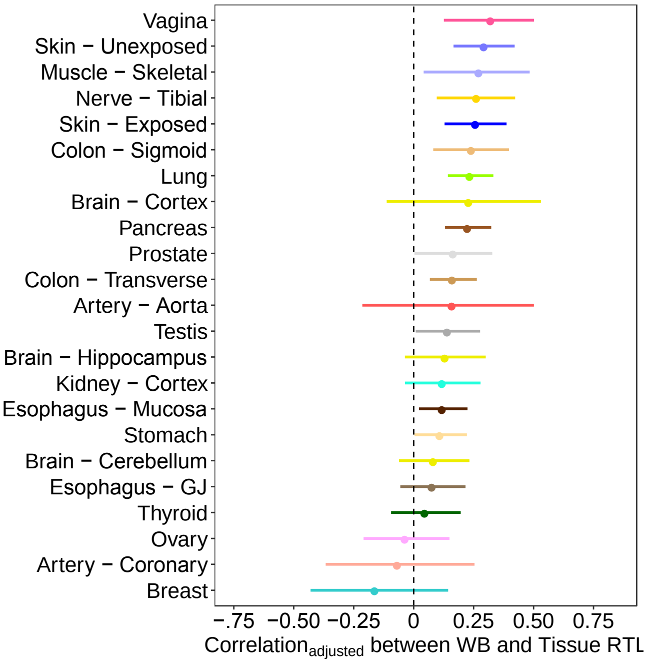


**Fig. S5.**

Forest plot of partial correlations between whole blood RTL and tissue-specific TL. Partial correlation adjusted for age, sex, race/ethnicity, BMI, and donor ischemic time. Black dashed line corresponds to $r_{partial}$=0.

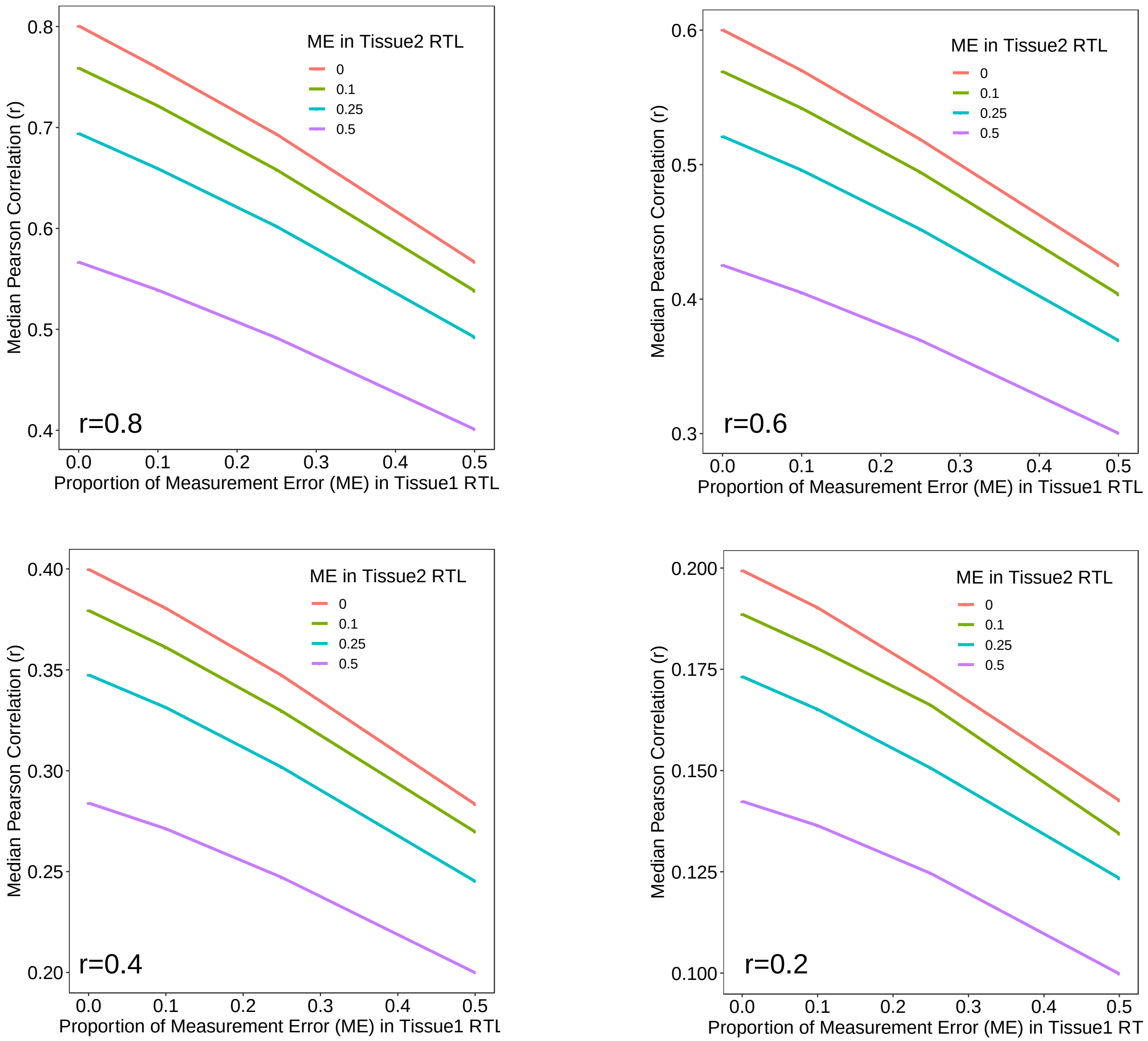


**Fig. S6.**

Results from analyses of simulated datasets examining effect of non-differential (random) measurement error on the observed Pearson correlation (r) between RTL measures taken from different two tissue types. Simulation sample size was 500, and we ran 1,000 simulations per scenario. The parameters for measurement error for tissue 1 RTL ($\sigma^2_{ux}$) and tissue 2 RTL ($\sigma^2_{uy}$) were varied to reflect the proportion of variation from 0 to 0.5 in RTL accounted for by measurement error. For more details, see **Methods.**

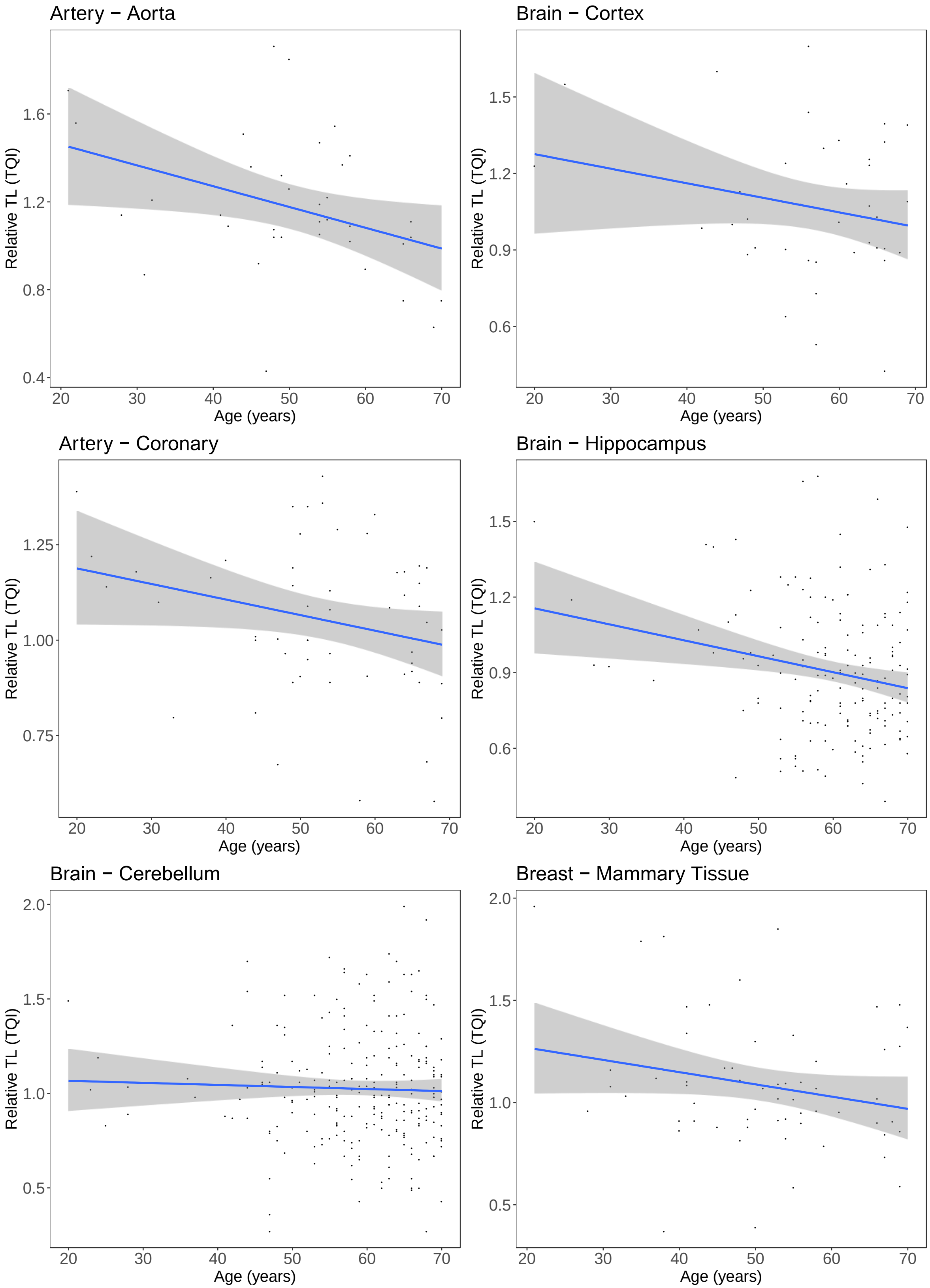


**Fig. S7.** Relationship between age and RTL for each tissue with ≥ 25 samples (24 tissue types). Blue line denotes line of best fit between RTL and age and gray 95% confidence interval bands.

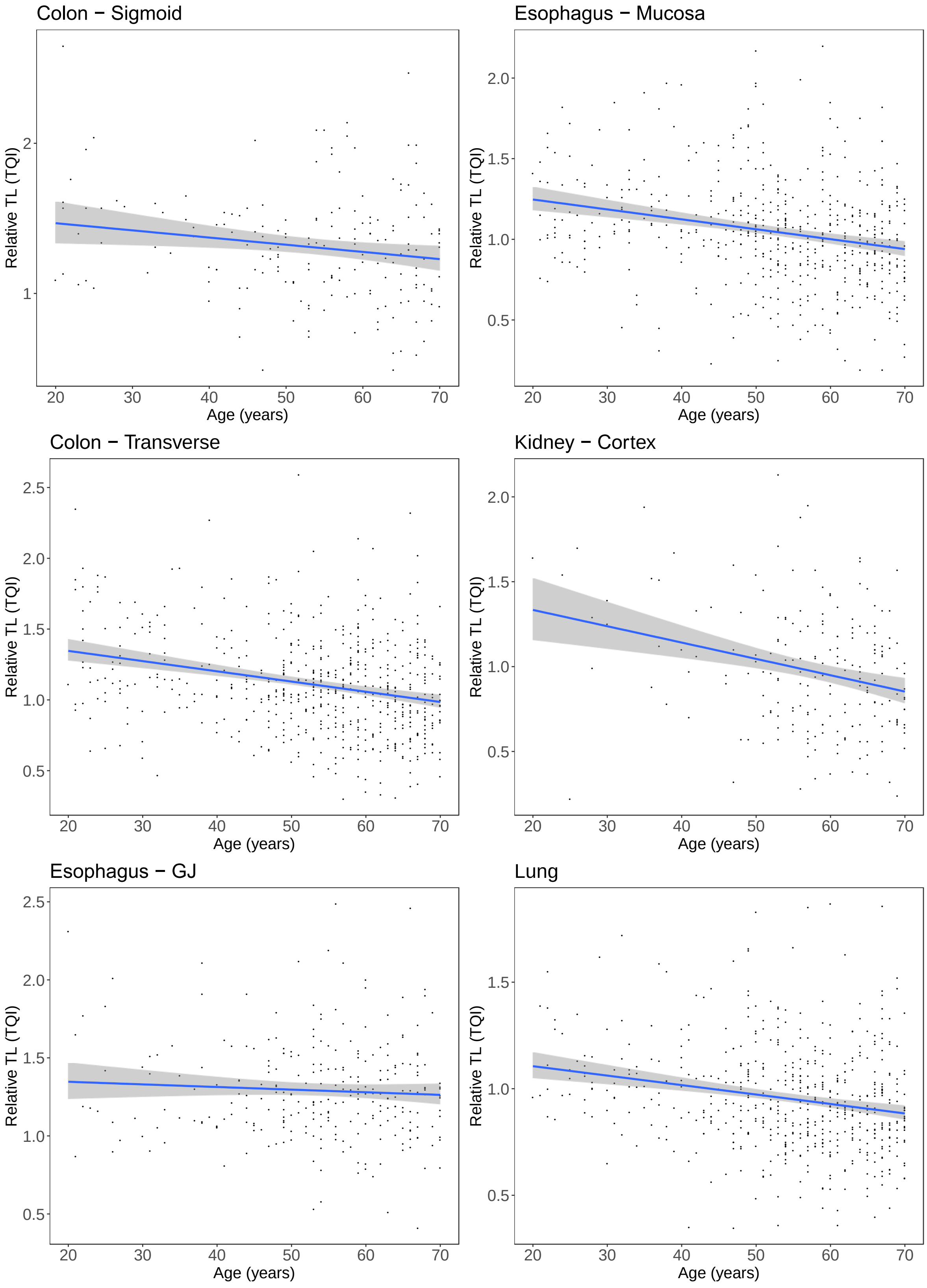


**Fig. S7 (Continued).**

Relationship between age and RTL for each tissue with ≥ 25 samples (n=24 tissues). Blue line denotes line of best fit between RTL and age and gray 95% confidence interval bands.

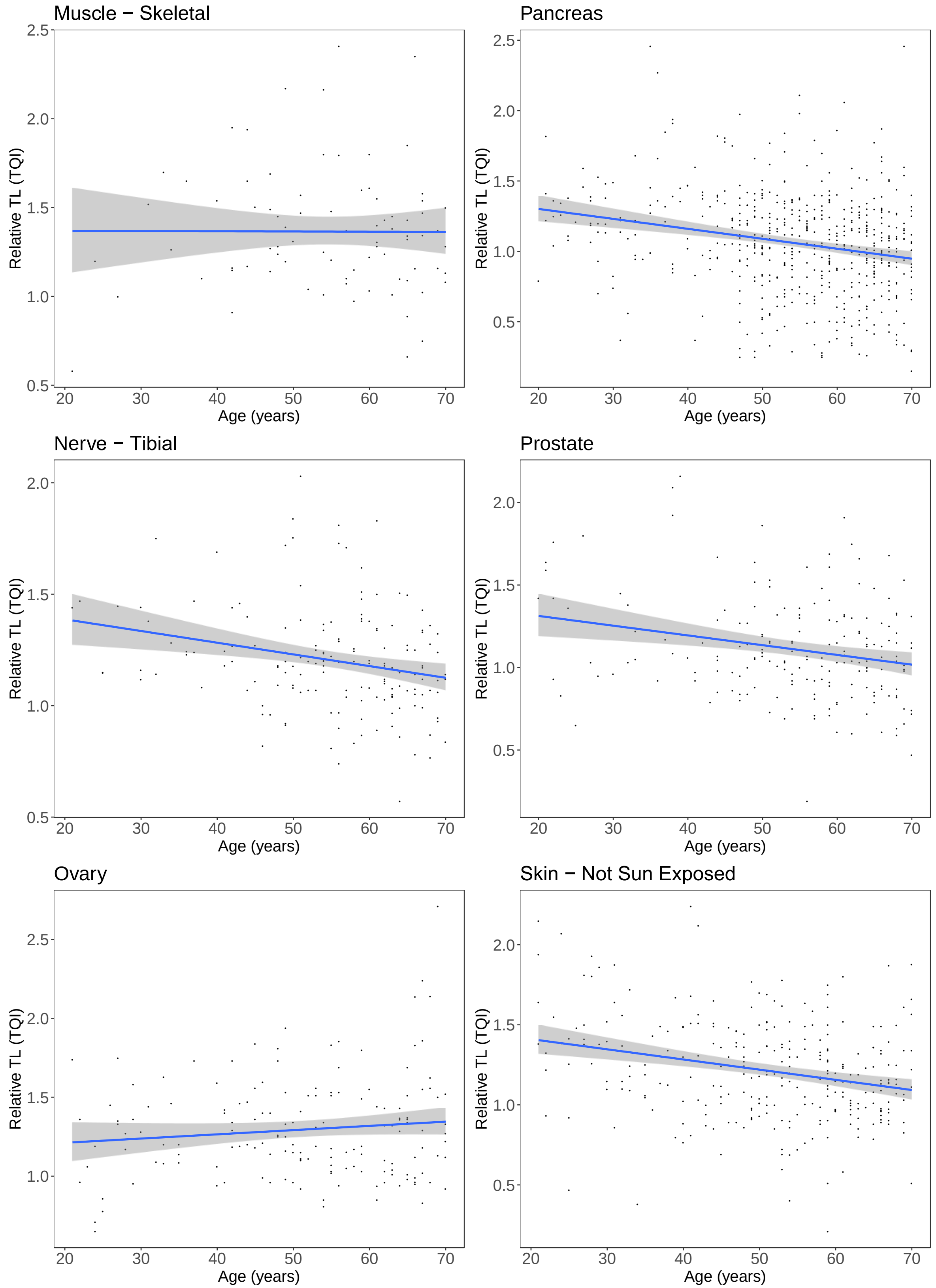


**Fig. S7 (Continued).**

Relationship between age and RTL for each tissue with ≥ 25 samples (n=24 tissues). Blue line denotes line of best fit between RTL and age and gray 95% confidence interval bands.

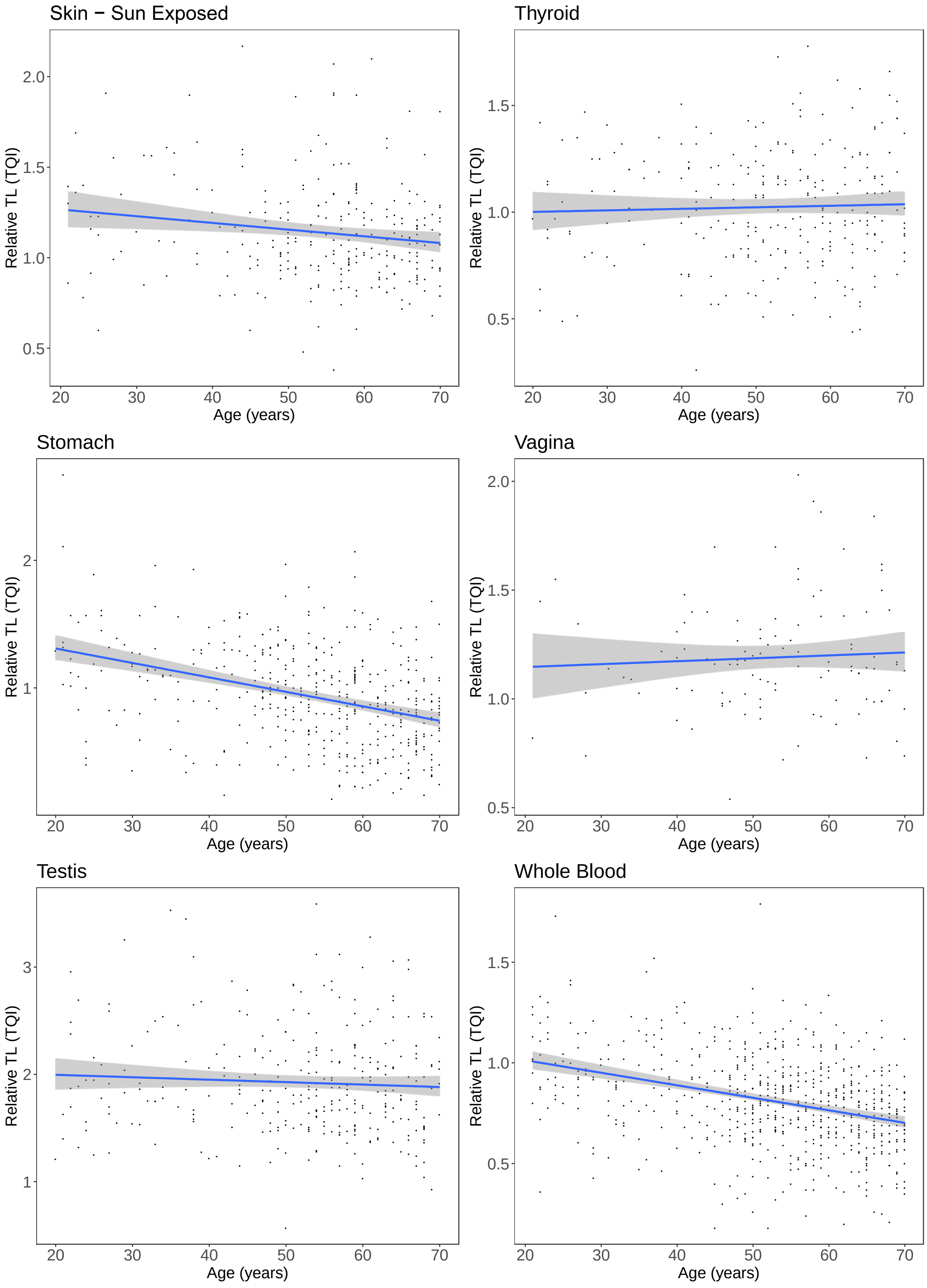


**Fig. S7 (Continued).**

Relationship between age and RTL for each tissue with ≥ 25 samples (n=24 tissues). Blue line denotes line of best fit between RTL and age and gray 95% confidence interval bands.

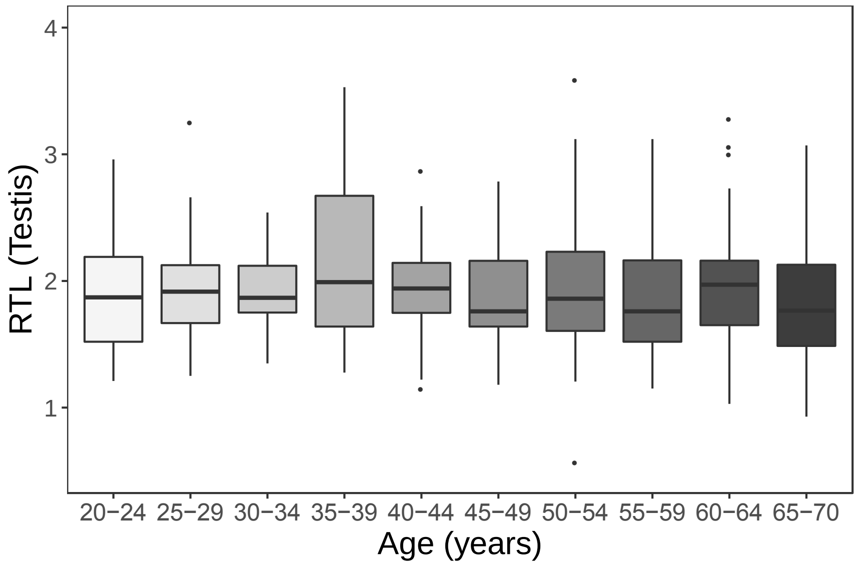


**Fig. S8.**

Testis RTL distribution across 5-year age groups (n=306).

A.

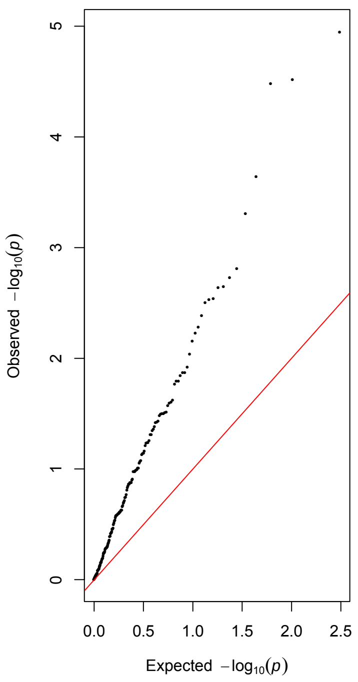


B.

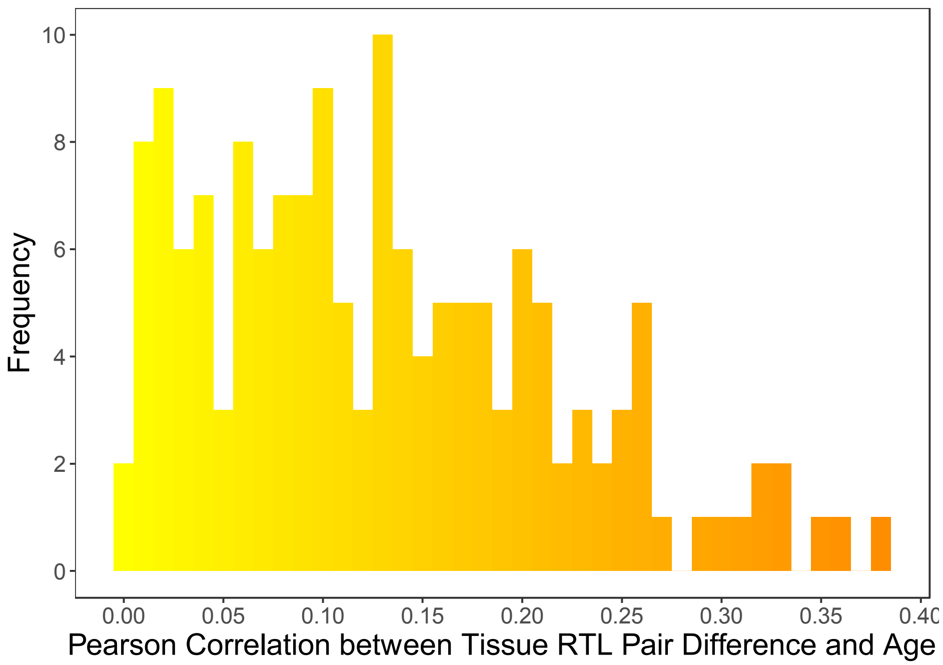


**Fig. S9.**

Tissue RTL pair differences and relationship with age (n=155 pairs). We calculated the difference in RTL between all pairs of tissue types (ΔRTL) available for each donor and restricted to tissue pairs with complete data for 50 ≥ donors. A) Q-Q plot of p-values from Pearson correlations between ΔRTL and age for all tissue pairs examined. B) Distribution of absolute Pearson correlations between ΔRTL and age for all tissue pairs examined.

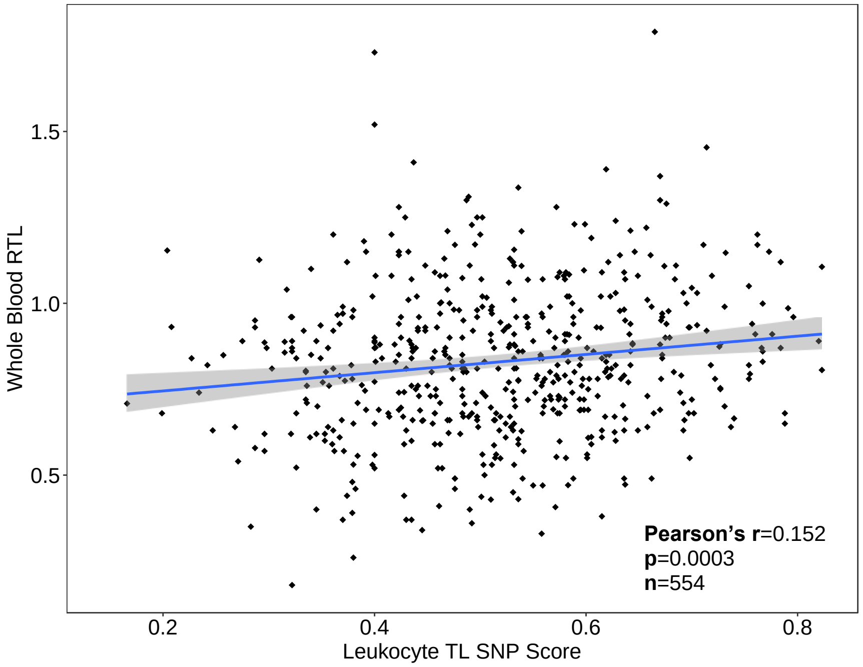


**Fig. S10.**

Correlation between polygenic SNP Score and RTL in whole blood (n=559).

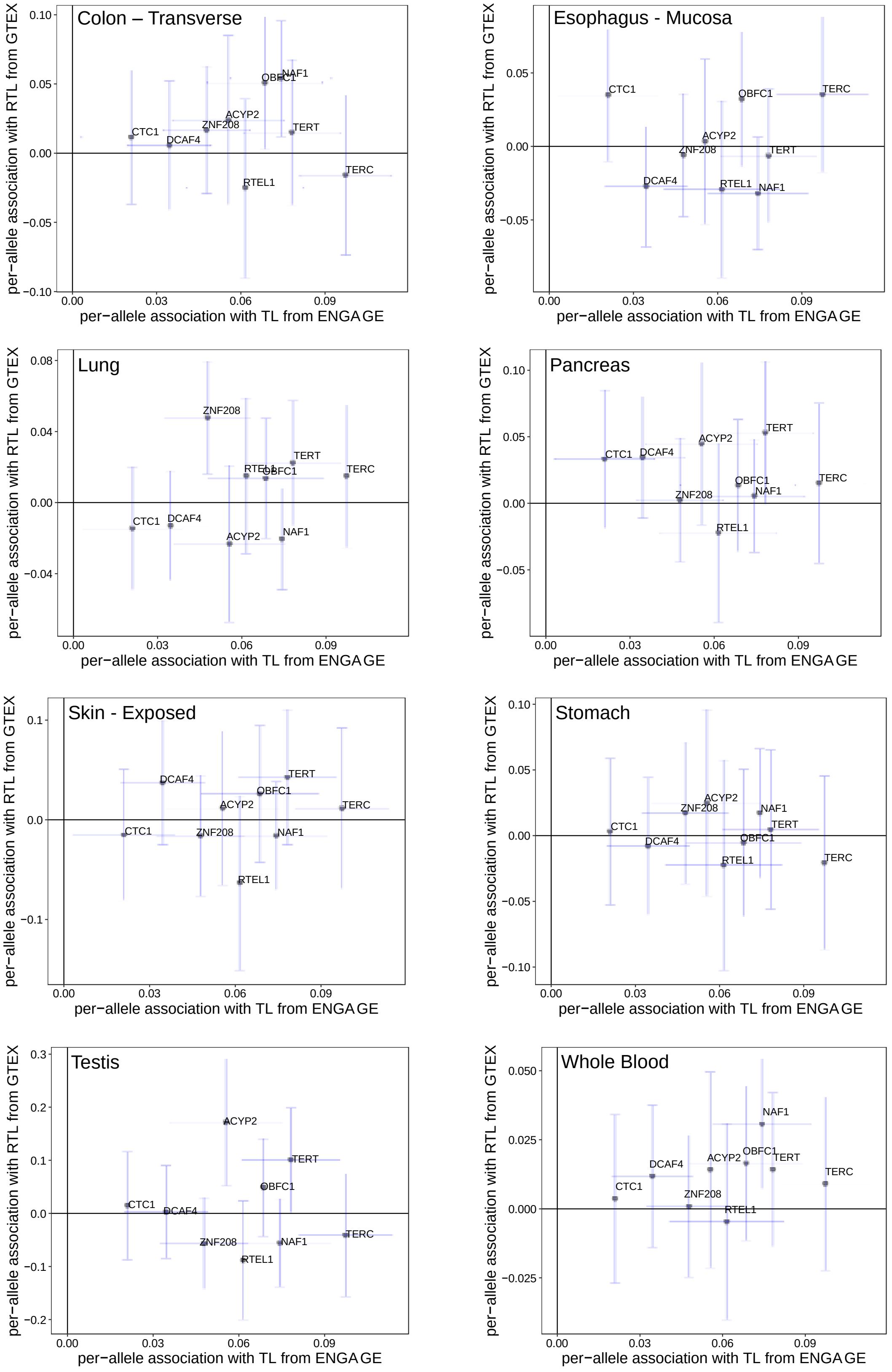


**Fig. S11.**

Relationship between TL-associated SNPs and leukocyte TL from ENGAGE (x-axis) and RTL from GTEx (tissues with ≥ 250 samples with RTL and genotype data).

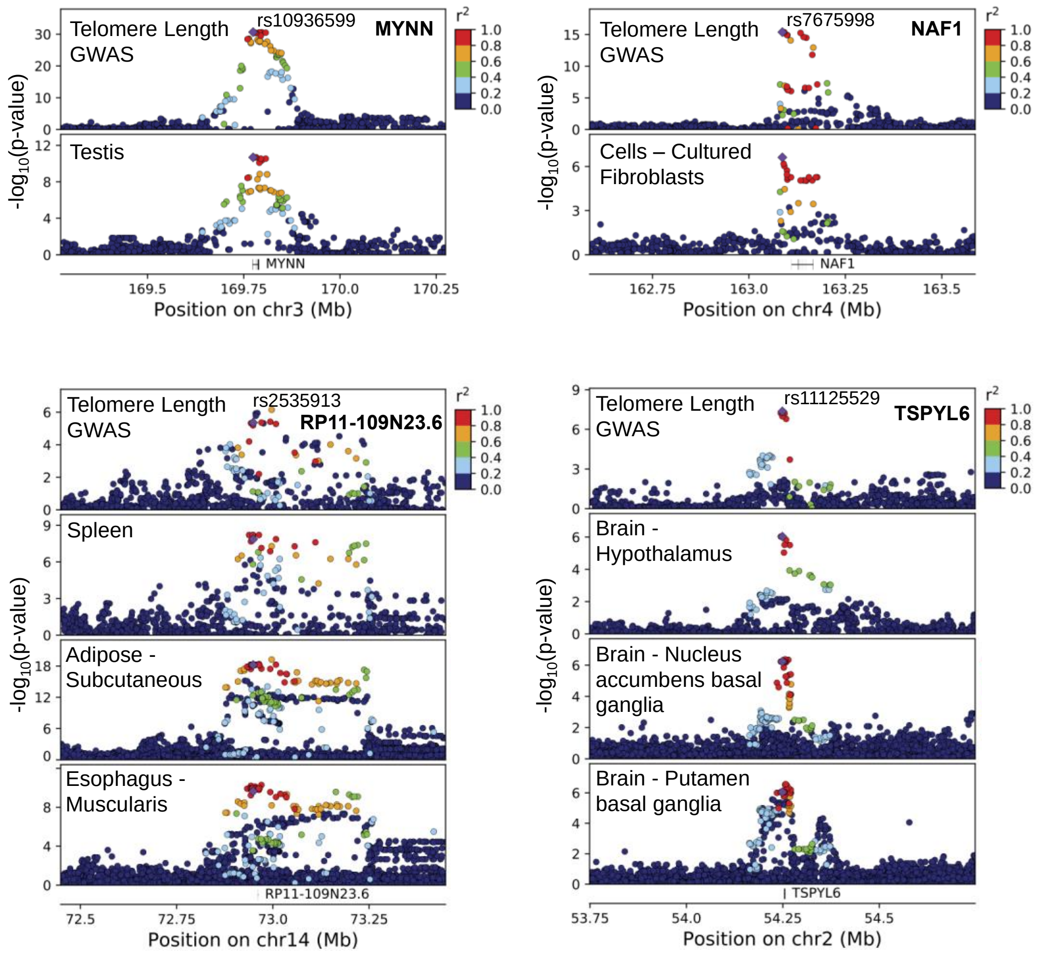


**Fig. S12.**

Association signals for leukocyte telomere length from prior GWAS (Codd, et al. 2013) co-localize with *cis*-eQTLs affecting expression of nearby genes human tissue types and cell lines (based on GTEx eQTL results).

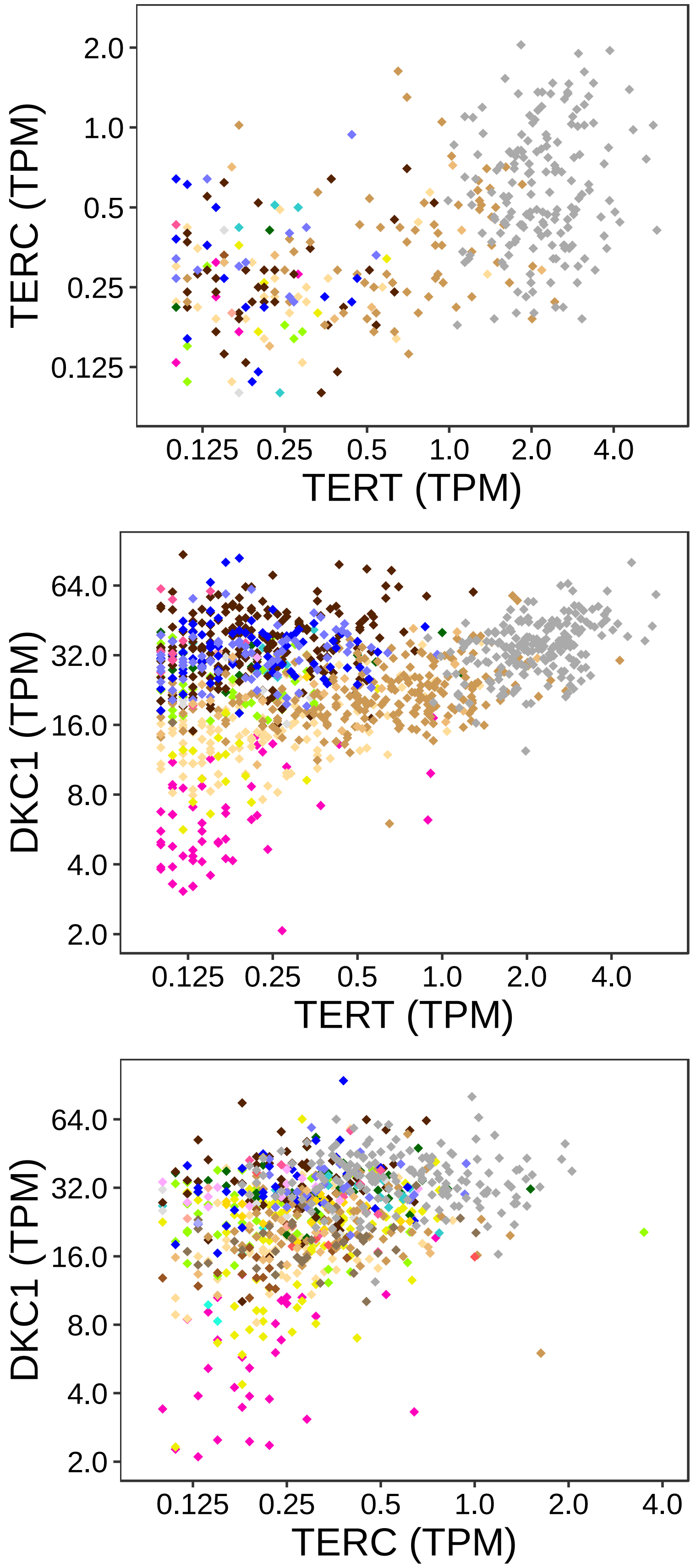


**Fig. S13.**

Relationship between *TERT*, *TERC*, and *DKC1* expression across all samples.

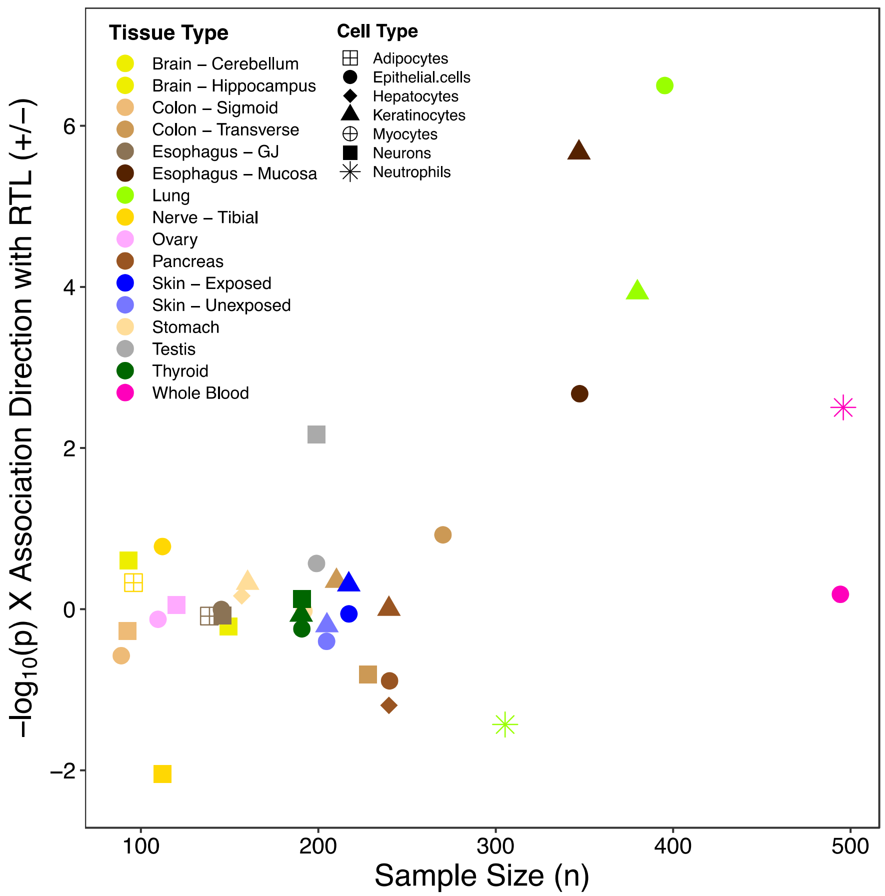


**Fig. S14.**

Selected GTEx cell type estimates are associated with TL in some tissue types. Reported direction of association and corresponding p-value between cell type enrichment scores (CTES) (estimated by xCell) and RTL adjusted for age and sex. Estimates reported for CTES within tissue types with ≥ 100 samples and detected in 75% or more samples and median CTES > 0.01 within a tissue type.

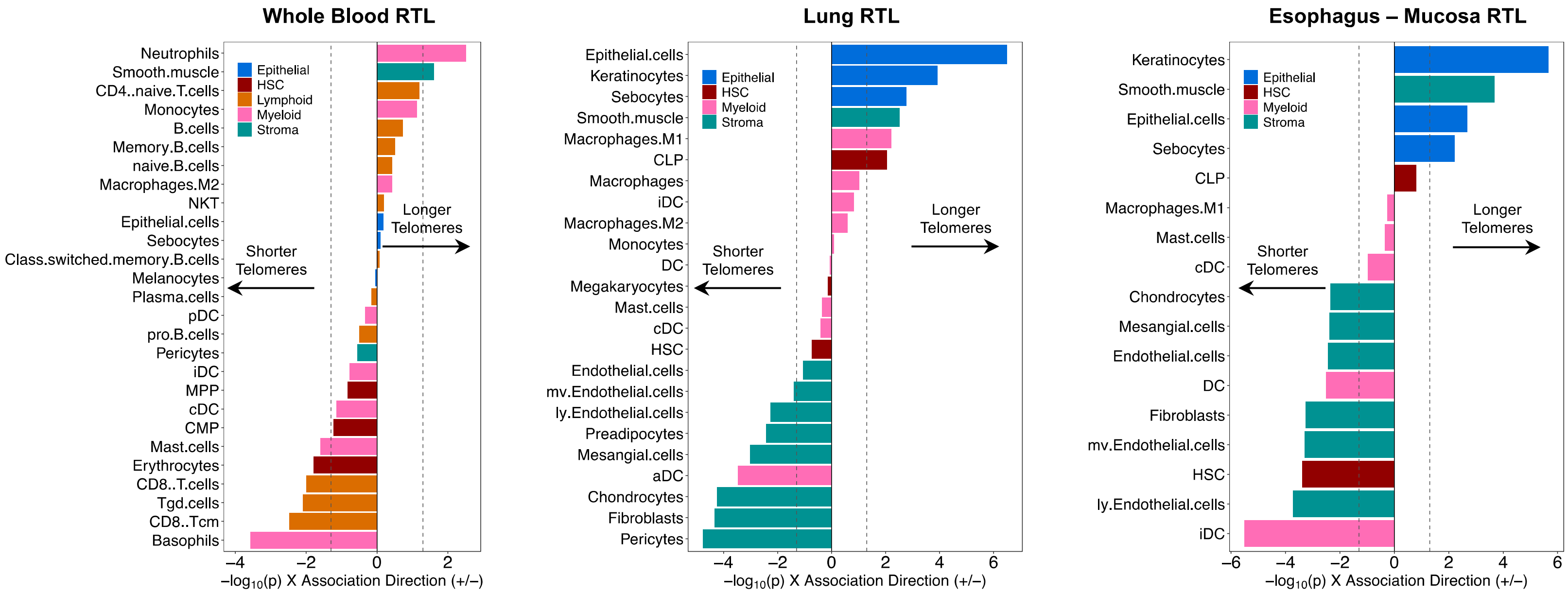


**Fig. S15.**

Cell type composition is associated with TL within tissue types. Reported direction of association and corresponding p-value between cell type enrichment scores (CTES) (estimated by xCell for whole blood, esophagus (mucosa), and lung samples) and RTL adjusted for age and sex. Estimates reported for CTES detected in 90% or more samples and median CTES > 0.01 within a tissue type. Colors correspond to cell type subgroup: hematopoietic stem cell (dark red), lymphoid (orange), myeloid (pink), stroma (teal), and epithelial (blue).

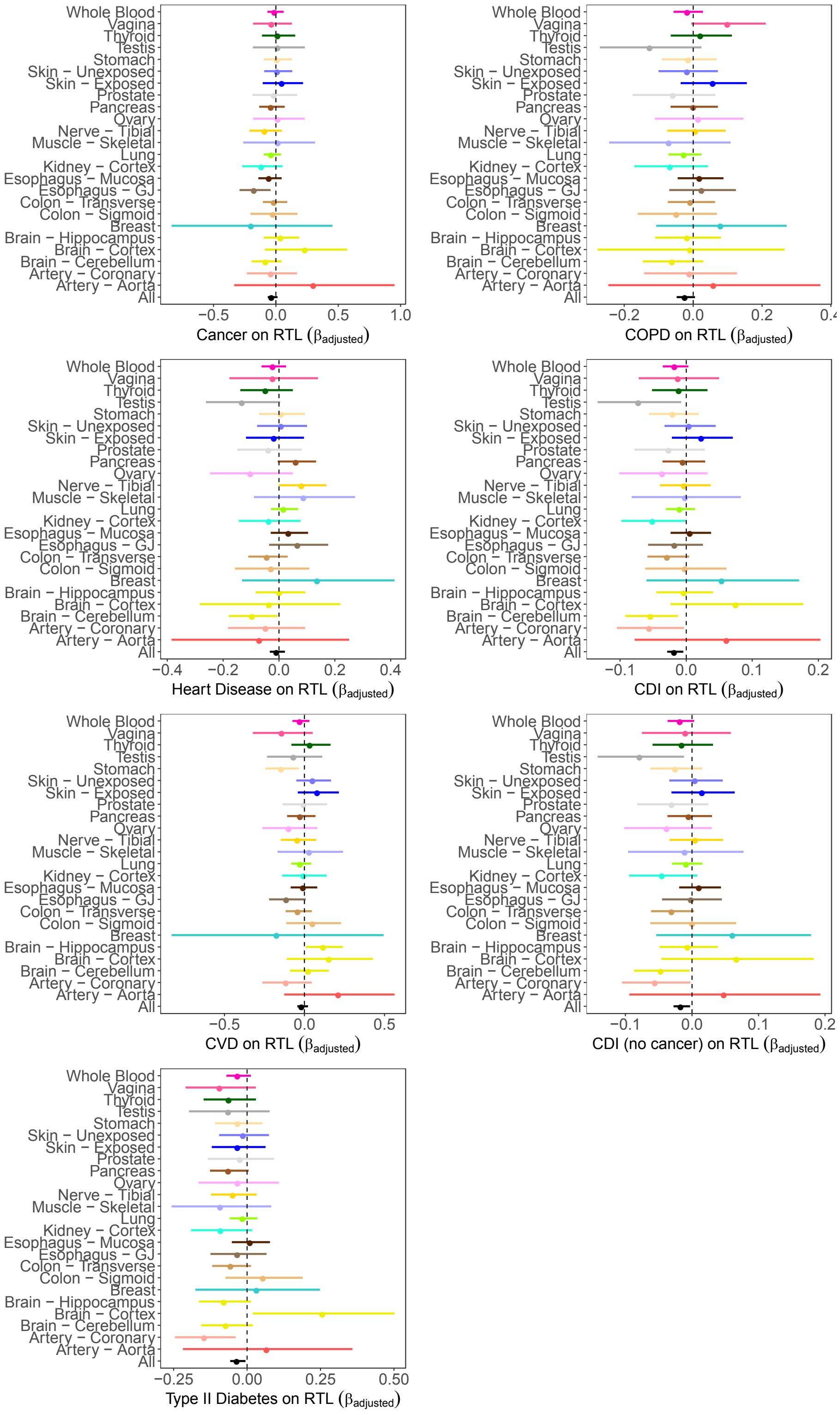


**Fig. S16.**

Shorter tissue RTL is associated with a history of chronic disease. Forest plot of associations (β) between five most common chronic diseases diagnosed among adults 20-70 years and chronic disease index. The prevalence of chronic disease was 7% for cancer (n=68 donors), 19% for COPD (n=178), 11% for CVD (n=106), 20% for HD (n=185), and 22% for T2D (n=209). Among the donors, 50% had no history of any chronic disease, and 30%, 14%, and 6% had a history of one, two, and three (or more) chronic diseases, respectively. For within tissue analyses, linear mixed models were adjusted for age, sex, race/ethnicity, BMI, donor ischemic time, and technical factors (DNA concentration and sample plate) as a random effect. For across tissue analysis, linear mixed models were adjusted for age, sex, race/ethnicity, BMI, donor ischemic time, technical factors, and random effect of tissue type and donor. Black dashed line corresponds to β=0.

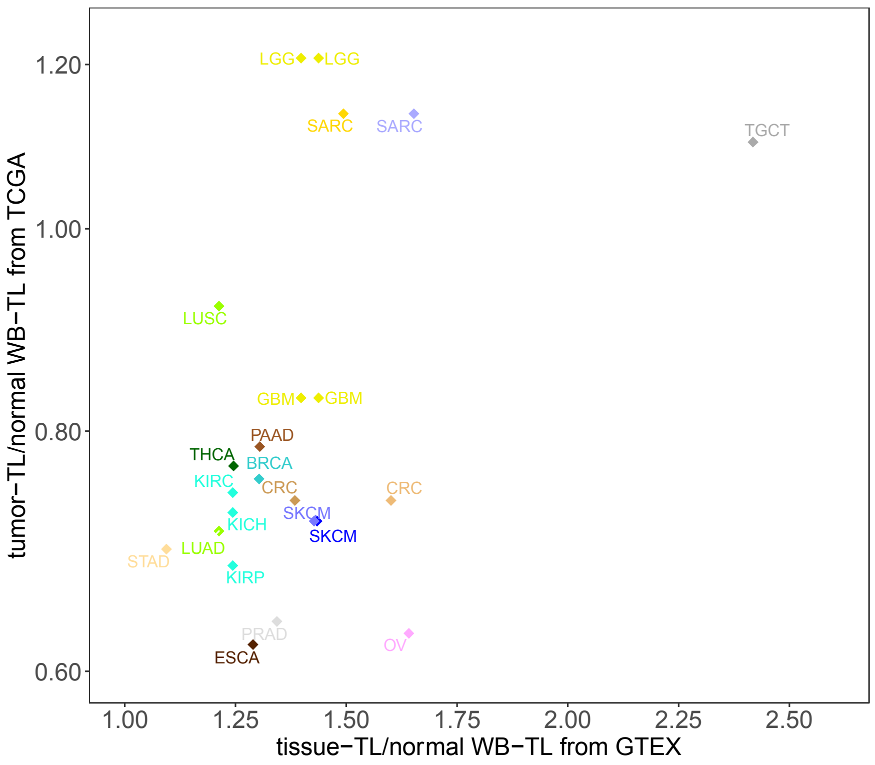


**Fig. S17.**

GTEx normal tissue TL and TCGA cancer tissue TL are positively correlated. For both GTEx and TCGA, the mean of the ratio of the tissue (or cancer) TL to the normal WB TL were utilized in the comparison. Abbreviations for cancers are BRCA (breast invasive carcinoma), CRC (colorectal carcinoma), ESCA (esophageal carcinoma), GBM (glioblastoma multiforme), KICH (kidney chromophobe), KIRC (kidney renal clear cell carcinoma), KIRP (kidney renal papillary cell carcinoma), LGG (low grade glioma), LUAD (lung adenocarcinoma), LUSC (lung squamous cell carcinoma), OV (ovarian serous cystadenocarcinoma), PAAD (pancreatic adenocarcinoma), PRAD (prostate adenocarcinoma), SARC (sarcoma), SKCM (skin cutaneous melanoma), STAD (stomach adenocarcinoma), THCA(thyroid carcinoma), and TGCT (testicular germ cell tumors ). Some tissues types and cancers are repeated if needed (see **Methods**).

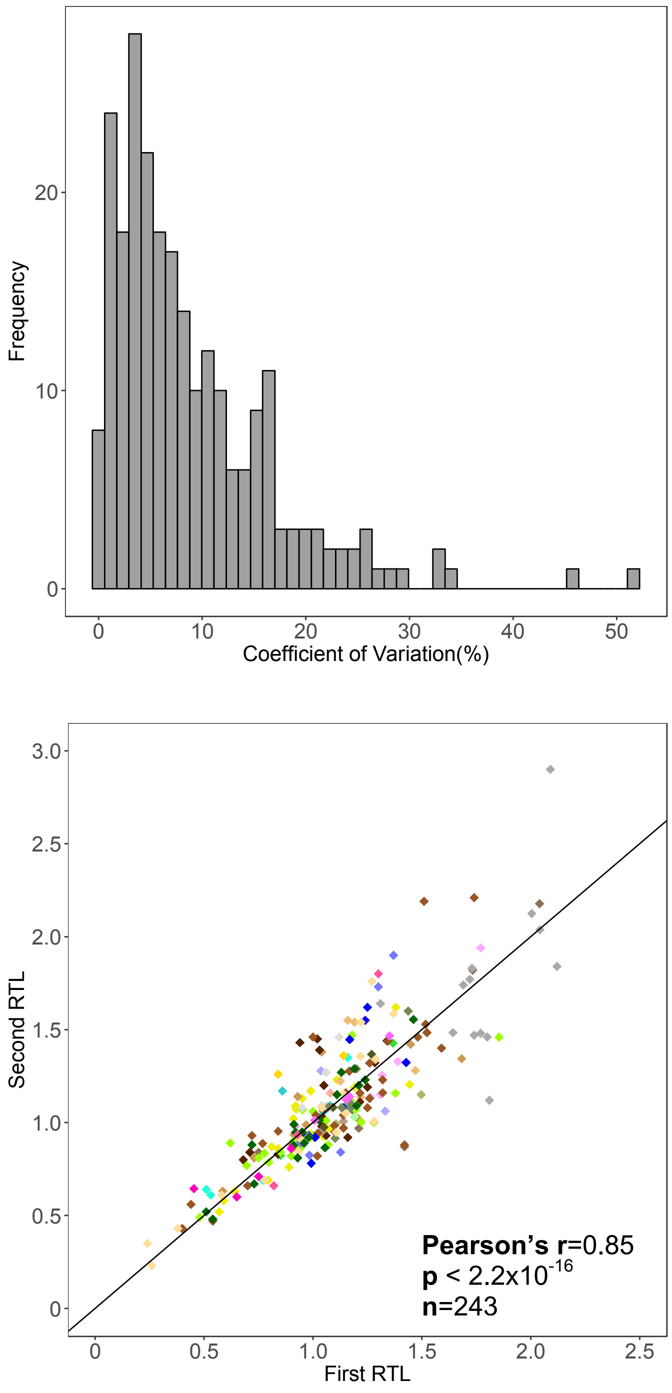


**Fig. S18.**

Assessment of repeated relative TL (RTL) measurements from GTEx samples (n=243). A) Distribution of coefficient of variation between each sample pair. B) First and second TL measurements for each sample pair. Black line reflects line for identical measurements (intercept=0, β=1).

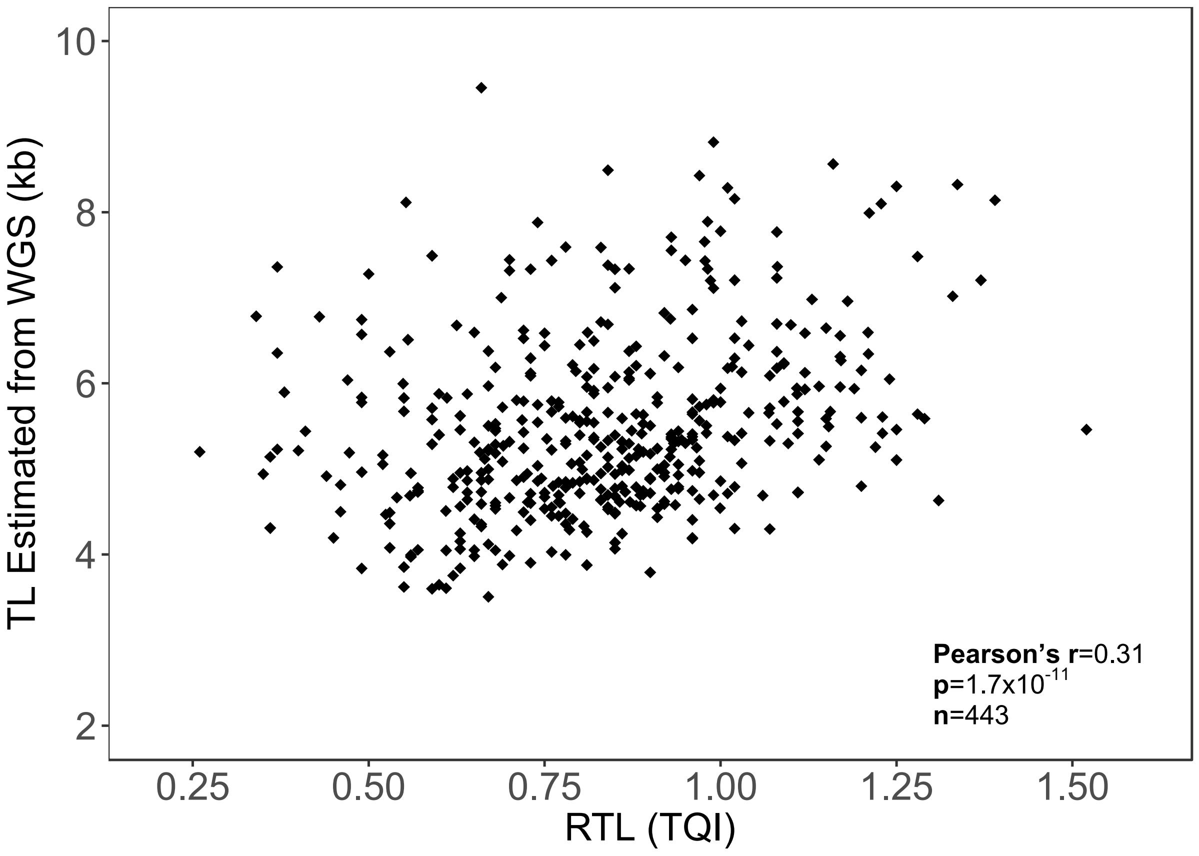


**Fig. S19.**

Comparison of RTL TL estimated from whole genome sequencing (WGS) of DNA from GTEx whole blood samples (n=443).

| Tissue | Color | n (attempted) | n (with RTL) | Missing | RTL mean (sd) | Age mean (sd) | Men (%) | African American (%) | White (%) | Organ Donor (%) |
|---|---|---|---|---|---|---|---|---|---|---|
| Artery - Aorta | | 43 | 36 | 7 (16.3%) | 1.17 (0.32) | 50.4 (12.2) | 63.9 | 8.3 | 91.7 | 58.3 |
| Artery - Coronary | | 63 | 55 | 8 (12.7%) | 1.05 (0.20) | 53.2 (12.6) | 67.3 | 12.7 | 87.3 | 50.9 |
| Brain - Cerebellum | | 250 | 241 | 9 (3.6%) | 1.02 (0.30) | 59.1 (9.3) | 71.0 | 9.5 | 89.2 | 0.0 |
| Brain - Cortex | | 49 | 39 | 10 (20.4%) | 1.07 (0.28) | 56.7 (11.1) | 53.8 | 5.1 | 92.3 | 0.0 |
| Brain - Hippocampus | | 188 | 160 | 28 (14.9%) | 0.90 (0.26) | 60.0 (8.9) | 72.5 | 9.4 | 90.6 | 0.0 |
| Breast - Mammary Tissue | | 66 | 61 | 5 (7.6%) | 1.08 (0.32) | 51.1 (11.7) | 24.6 | 14.8 | 85.2 | 59.0 |
| Colon - Sigmoid | | 190 | 183 | 7 (3.7%) | 1.31 (0.36) | 53.4 (13.3) | 58.5 | 9.3 | 90.2 | 43.7 |
| Colon - Transverse | | 614 | 568 | 46 (7.5%) | 1.11 (0.35) | 53.1 (13.0) | 67.8 | 12.0 | 85.2 | 45.2 |
| Esophagus - GJ | | 281 | 267 | 14 (5.0%) | 1.29 (0.32) | 53.6 (11.9) | 62.9 | 10.1 | 87.6 | 47.2 |
| Esophagus - Mucosa | | 588 | 528 | 60 (10.2%) | 1.05 (0.33) | 52.4 (13.1) | 67.2 | 11.6 | 86.4 | 47.3 |
| Kidney - Cortex | | 212 | 202 | 10 (4.7%) | 0.97 (0.36) | 57.7 (10.2) | 66.3 | 7.9 | 90.6 | 6.4 |
| Lung | | 639 | 556 | 83 (13.0%) | 0.95 (0.25) | 54.2 (11.9) | 67.4 | 10.8 | 87.1 | 40.5 |
| Muscle - Skeletal | | 86 | 80 | 6 (7.0%) | 1.36 (0.35) | 54.5 (11.7) | 57.5 | 18.8 | 81.3 | 35.0 |
| Nerve - Tibial | | 193 | 162 | 31 (16.1%) | 1.20 (0.24) | 54.9 (11.2) | 66.0 | 9.3 | 88.3 | 43.2 |
| Ovary | | 166 | 155 | 11 (6.6%) | 1.29 (0.32) | 51.1 (13.7) | 0.0 | 14.8 | 83.2 | 52.3 |
| Pancreas | | 620 | 546 | 74 (11.9%) | 1.06 (0.36) | 54.4 (11.7) | 67.2 | 10.8 | 86.4 | 43.8 |
| Prostate | | 201 | 188 | 13 (6.5%) | 1.11 (0.31) | 54.4 (12.5) | 100.0 | 10.6 | 87.8 | 33.5 |
| Skin - Not Sun Exposed | | 280 | 263 | 17 (6.1%) | 1.14 (0.29) | 54.3 (12.3) | 68.8 | 10.3 | 87.1 | 41.4 |
| Skin - Sun Exposed | | 367 | 286 | 81 (22.1%) | 1.21 (0.32) | 51.2 (12.9) | 65.4 | 12.9 | 83.6 | 45.8 |
| Stomach | | 471 | 420 | 51 (10.8%) | 0.93 (0.39) | 53.1 (12.7) | 64.5 | 9.3 | 89.5 | 46.7 |
| Testis | | 337 | 306 | 31 (9.2%) | 1.92 (0.49) | 51.8 (13.1) | 100.0 | 10.8 | 86.6 | 44.4 |
| Thyroid | | 267 | 255 | 12 (4.5%) | 1.02 (0.27) | 51.9 (12.9) | 56.5 | 10.6 | 87.8 | 51.4 |
| Vagina | | 113 | 108 | 5 (4.4%) | 1.19 (0.27) | 52.7 (11.7) | 0.0 | 14.8 | 82.4 | 48.1 |
| Whole Blood | | 841 | 637 | 204 (24.3%) | 0.81 (0.23) | 52.5 (12.8) | 65.5 | 14.8 | 82.9 | 46.0 |
| All Samples | | 7227 | 6391 | 836 (11.6%) | 1.10 (0.39) | | | | | |
| All Donors | | 962 | 952 | 10 (1.0%) | | 52.8 (12.9) | 66.6 | 12.7 | 84.9 | 44.7 |

**Table S1** Summary of participants and tissue samples with TL measurements (restricting to tissue types with ≥ 25 samples measured).

| | | All Tissues | | Excluding Testis | |
|---|---|---|---|---|---|
| **Factor** | **Model** | **PVE** | **p (LRT)** | **PVE** | **p (LRT)** |
| All Fixed Effects | ~ Age + Sex + Race + BMI + Ischemic Time + Plate + (1\|Tissue) + (1\|Participant) | 16.7% | $<2.2 \times 10^{-16}$ | 20.4% | $<2.2 \times 10^{-16}$ |
| Age | ~ Age + (1\|Tissue) + (1\|Participant) | 3.3% | $<2.2 \times 10^{-16}$ | 4.4% | $<2.2 \times 10^{-16}$ |
| Sex | ~ Sex + (1\|Tissue) + (1\|Participant) | 0.0% | $2.8 \times 10^{-1}$ | 0.0% | $2.7 \times 10^{-1}$ |
| Race/Ethnicity | ~ Race + (1\|Tissue) + (1\|Participant) | 0.6% | $3.0 \times 10^{-5}$ | 0.7% | $2.0 \times 10^{-5}$ |
| BMI | ~ BMI + (1\|Tissue) + (1\|Participant) | 0.4% | $9.2 \times 10^{-5}$ | 0.5% | $1.1 \times 10^{-4}$ |
| Donor Ischemic Time | ~ Ischemic Time + (1\|Tissue) + (1\|Participant) | 3.7% | $<2.2 \times 10^{-16}$ | 5.3% | $<2.2 \times 10^{-16}$ |
| Plate | ~ Plate + (1\|Tissue) + (1\|Participant) | 11.2% | $<2.2 \times 10^{-16}$ | 12.8% | $<2.2 \times 10^{-16}$ |
| Tissue (from random effect) | ~ Age + Sex + Race + BMI + Ischemic Time + Plate + (1\|Tissue) + (1\|Participant) | 24.3% | $<2.2 \times 10^{-16}$ | 11.5% | $<2.2 \times 10^{-16}$ |
| Individual (from random effect) | ~ Age + Sex + Race + BMI + Ischemic Time + Plate + (1\|Tissue) + (1\|Participant) | 8.7% | $<2.2 \times 10^{-16}$ | 11.2% | $<2.2 \times 10^{-16}$ |

**Table S2**

Estimates of variation of telomere length explained by each fixed effect. Linear mixed models were utilized with random effects for tissue type and donor in all models except when noted. For fixed effects, we extracted estimates from marginal $R^2$ using approach of Nagawa et al (2013) that represents the marginal percent variation explained (PVE) and *p*-value was obtained from likelihood ratio test comparing model with the fixed effect and without the fixed effect.

| | Males | | Females | | Sex | | | |
|---|---|---|---|---|---|---|---|---|
| **Tissue** | **n** | **mean (sd)** | **n** | **mean (sd)** | $\beta_{unadjusted}$ | $p_{unadjusted}$ | $\beta_{adjusted}$ | $p_{adjusted}$ |
| Brain - Cerebellum | 171 | 1.02 (0.30) | 70 | 1.04 (0.31) | 0.017 (0.043) | 0.6824 | 0.008 (0.042) | 0.8508 |
| Brain - Hippocampus | 116 | 0.91 (0.26) | 44 | 0.88 (0.25) | -0.030 (0.046) | 0.5168 | -0.032 (0.045) | 0.4859 |
| Colon - Sigmoid | 107 | 1.32 (0.38) | 76 | 1.28 (0.34) | -0.040 (0.054) | 0.4643 | -0.037 (0.051) | 0.4547 |
| Colon - Transverse | 385 | 1.10 (0.35) | 183 | 1.12 (0.35) | 0.012 (0.031) | 0.6973 | -0.023 (0.029) | 0.4134 |
| Esophagus - GJ | 168 | 1.30 (0.34) | 99 | 1.28 (0.28) | -0.020 (0.041) | 0.6200 | -0.027 (0.038) | 0.4764 |
| Esophagus - Mucosa | 355 | 1.05 (0.33) | 173 | 1.05 (0.32) | -0.003 (0.030) | 0.9324 | -0.028 (0.027) | 0.2930 |
| Kidney - Cortex | 134 | 1.00 (0.38) | 68 | 0.92 (0.30) | -0.072 (0.053) | 0.1781 | -0.077 (0.048) | 0.1033 |
| Lung | 375 | 0.94 (0.24) | 181 | 0.98 (0.25) | 0.035 (0.022) | 0.1141 | 0.016 (0.020) | 0.4396 |
| Muscle - Skeletal | 46 | 1.42 (0.37) | 34 | 1.29 (0.30) | -0.131 (0.077) | 0.0935 | **-0.170 (0.077)** | **0.0244** |
| Nerve - Tibial | 107 | 1.22 (0.24) | 55 | 1.16 (0.23) | -0.061 (0.039) | 0.1168 | -0.058 (0.037) | 0.1116 |
| Pancreas | 367 | 1.03 (0.36) | 179 | 1.12 (0.35) | **0.096 (0.033)** | **0.0034** | **0.070 (0.029)** | **0.0162** |
| Skin - Not Sun Exposed | 181 | 1.13 (0.29) | 82 | 1.16 (0.29) | 0.026 (0.038) | 0.4961 | 0.039 (0.037) | 0.2836 |
| Skin - Sun Exposed | 187 | 1.22 (0.33) | 99 | 1.20 (0.29) | -0.015 (0.039) | 0.6737 | -0.042 (0.037) | 0.2652 |
| Stomach | 271 | 0.92 (0.40) | 149 | 0.96 (0.38) | 0.037 (0.040) | 0.3501 | -0.019 (0.033) | 0.5646 |
| Thyroid | 144 | 1.03 (0.27) | 111 | 1.02 (0.27) | -0.007 (0.034) | 0.8411 | 0.005 (0.034) | 0.9034 |
| Whole Blood | 417 | 0.81 (0.23) | 220 | 0.82 (0.23) | 0.007 (0.019) | 0.7109 | -0.005 (0.018) | 0.7940 |

**Table S3**

Association between sex and RTL within tissues (non-reproductive tissues and n ≥ 75). Linear mixed models were used to estimate the association (β) between sex and RTL within each tissue, and we obtained *p*-value from likelihood ratio test comparing model with and without sex. Estimates with *p*-values < 0.05 are shown in bold. Adjusted linear mixed models included race/ethnicity, age, donor ischemic time, BMI, and technical factors (DNA concentration and sample plate) as a random effect.

| | European Ancestry | | African Ancestry | | Reported Association between Ancestry (EA v.s. AA) and RTL within Tissue | | | |
|---|---|---|---|---|---|---|---|---|
| **Tissue** | **n** | **mean (sd)** | **n** | **mean (sd)** | $\beta_{unadjusted}$ | $p_{unadjusted}$ | $\beta_{adjusted}$ | $p_{adjusted}$ |
| Brain - Cerebellum | 170 | 1.02 (0.30) | 18 | 1.19 (0.28) | 0.171 (0.073) | **0.0207** | 0.161 (0.073) | **0.0268** |
| Brain - Hippocampus | 121 | 0.90 (0.25) | 14 | 0.95 (0.36) | 0.053 (0.073) | 0.4714 | 0.042 (0.073) | 0.5589 |
| Colon - Sigmoid | 147 | 1.33 (0.38) | 15 | 1.32 (0.24) | -0.011 (0.099) | 0.9121 | -0.006 (0.094) | 0.9476 |
| Colon - Transverse | 411 | 1.10 (0.36) | 57 | 1.19 (0.34) | 0.089 (0.051) | 0.0798 | 0.047 (0.046) | 0.3039 |
| Esophagus - GJ | 206 | 1.30 (0.33) | 21 | 1.39 (0.32) | 0.083 (0.076) | 0.2753 | 0.064 (0.072) | 0.3653 |
| Esophagus - Mucosa | 401 | 1.03 (0.32) | 53 | 1.17 (0.36) | 0.143 (0.047) | **0.0025** | 0.082 (0.042) | 0.0514 |
| Kidney - Cortex | 157 | 0.98 (0.37) | 16 | 1.14 (0.31) | 0.163 (0.095) | 0.0885 | 0.136 (0.089) | 0.1224 |
| Lung | 409 | 0.96 (0.24) | 51 | 1.06 (0.26) | 0.103 (0.036) | **0.0042** | 0.078 (0.033) | **0.0174** |
| Nerve - Tibial | 122 | 1.20 (0.23) | 13 | 1.30 (0.37) | 0.096 (0.072) | 0.1830 | 0.087 (0.067) | 0.1916 |
| Ovary | 113 | 1.29 (0.33) | 17 | 1.29 (0.32) | 0.009 (0.086) | 0.9183 | -0.003 (0.084) | 0.9697 |
| Pancreas | 394 | 1.06 (0.36) | 48 | 1.12 (0.37) | 0.062 (0.055) | 0.2610 | 0.010 (0.048) | 0.8448 |
| Prostate | 144 | 1.11 (0.30) | 17 | 1.26 (0.40) | 0.152 (0.080) | 0.0543 | 0.163 (0.076) | **0.0316** |
| Skin - Not Sun Exposed | 216 | 1.22 (0.32) | 31 | 1.24 (0.34) | 0.019 (0.062) | 0.7231 | -0.007 (0.059) | 0.9168 |
| Skin - Sun Exposed | 199 | 1.13 (0.29) | 23 | 1.22 (0.36) | 0.096 (0.066) | 0.1455 | 0.070 (0.062) | 0.2556 |
| Stomach | 324 | 0.94 (0.40) | 35 | 1.04 (0.39) | 0.105 (0.072) | 0.1433 | 0.034 (0.060) | 0.5683 |
| Testis | 233 | 1.94 (0.51) | 28 | 1.98 (0.42) | 0.045 (0.100) | 0.6526 | 0.046 (0.096) | 0.6284 |
| Thyroid | 193 | 1.01 (0.28) | 23 | 1.15 (0.28) | 0.138 (0.061) | **0.0246** | 0.144 (0.061) | **0.0181** |
| Vagina | 80 | 1.20 (0.27) | 13 | 1.30 (0.26) | 0.103 (0.080) | 0.1979 | 0.097 (0.082) | 0.2231 |
| Whole Blood | 462 | 0.81 (0.21) | 78 | 0.90 (0.24) | 0.088 (0.026) | **0.0010** | 0.067 (0.024) | **0.0054** |
| All Tissues | 4784 | 1.10 (0.40) | 622 | 1.18 (0.39) | 0.095 (0.022) | **$1.1 \times 10^{-5}$** | 0.048 (0.018) | **0.0068** |

**Table S4**
Association between genetically-determined ancestry and RTL within tissue types ($n \geq 75$ samples). Linear mixed models were used to estimate the association ($\beta$) between ancestry and RTL within each tissue type and obtained *p*-values from likelihood ratio test comparing model with and without ancestry. Estimates with *p*-values <0.05 are shown in bold. Adjusted linear mixed models included sex, age, donor ischemic time, BMI, and technical factors (DNA concentration and sample plate) as a random effect. EA: European Ancestry; AA: African Ancestry.

| | All | | | | Males | | | | Females | | | | |
|---|---|---|---|---|---|---|---|---|---|---|---|---|---|
| Tissue | n | r | $\beta_{adjusted}$ | $p_{adjusted}$ | n | r | $\beta_{adjusted}$ | $p_{adjusted}$ | n | r | $\beta_{adjusted}$ | $p_{adjusted}$ | p (interaction) |
| Brain - Cerebellum | 241 | -0.034 | 0.000 (0.002) | $9.3x10^{-1}$ | 171 | 0.006 | 0.001 (0.003) | $6.6x10^{-1}$ | 70 | -0.106 | -0.001 (0.004) | $7.1x10^{-1}$ | 0.3444 |
| Brain - Hippocampus | 160 | -0.219 | -0.006 (0.002) | **$8.5x10^{-3}$** | 116 | -0.304 | -0.009 (0.003) | **$5.1x10^{-4}$** | 44 | 0.041 | 0.001 (0.005) | $7.5x10^{-1}$ | **0.0394** |
| Colon - Sigmoid | 183 | -0.177 | -0.003 (0.002) | $1.8x10^{-1}$ | 107 | -0.156 | -0.003 (0.003) | $2.7x10^{-1}$ | 76 | -0.207 | -0.003 (0.003) | $3.1x10^{-1}$ | 0.6380 |
| Colon - Transverse | 568 | -0.269 | -0.005 (0.001) | **$1.1x10^{-6}$** | 385 | -0.347 | -0.007 (0.001) | **$3.8x10^{-8}$** | 183 | -0.115 | -0.001 (0.002) | $5.7x10^{-1}$ | **0.0142** |
| Esophagus - GJ | 267 | -0.062 | -0.002 (0.002) | $1.5x10^{-1}$ | 168 | -0.109 | -0.003 (0.002) | $2.5x10^{-1}$ | 99 | 0.032 | 0.001 (0.003) | $7.9x10^{-1}$ | 0.2569 |
| Esophagus - Mucosa | 528 | -0.247 | -0.003 (0.001) | **$7.9x10^{-4}$** | 355 | -0.288 | -0.004 (0.001) | **$8.4x10^{-4}$** | 173 | -0.157 | -0.002 (0.002) | $4.1x10^{-1}$ | 0.1801 |
| Kidney - Cortex | 202 | -0.275 | -0.010 (0.002) | **$2.2x10^{-5}$** | 134 | -0.275 | -0.010 (0.003) | **$1.4x10^{-3}$** | 68 | -0.284 | -0.008 (0.003) | **$1.1x10^{-2}$** | 0.6414 |
| Lung | 556 | -0.215 | -0.003 (0.001) | **$8.6x10^{-4}$** | 375 | -0.275 | -0.004 (0.001) | **$1.3x10^{-4}$** | 181 | -0.096 | -0.001 (0.001) | $6.0x10^{-1}$ | **0.0430** |
| Muscle - Skeletal | 80 | -0.003 | 0.003 (0.004) | $4.7x10^{-1}$ | 46 | -0.078 | 0.001 (0.005) | $9.2x10^{-1}$ | 34 | 0.171 | 0.008 (0.006) | $1.8x10^{-1}$ | 0.2849 |
| Nerve - Tibial | 162 | -0.250 | -0.004 (0.002) | **$2.4x10^{-2}$** | 107 | -0.231 | -0.004 (0.002) | $5.4x10^{-2}$ | 55 | -0.299 | -0.007 (0.003) | **$3.2x10^{-2}$** | 0.5777 |
| Ovary | | | | | | | | | 155 | 0.115 | 0.002 (0.002) | $3.7x10^{-1}$ | |
| Pancreas | 546 | -0.229 | -0.003 (0.001) | **$2.8x10^{-2}$** | 367 | -0.279 | -0.003 (0.002) | **$4.1x10^{-2}$** | 179 | -0.132 | -0.002 (0.002) | $4.5x10^{-1}$ | 0.2993 |
| Prostate | | | | | 188 | -0.238 | -0.004 (0.002) | **$2.8x10^{-2}$** | | | | | |
| Skin - Not Sun Exposed | 263 | -0.159 | -0.003 (0.001) | **$2.0x10^{-2}$** | 181 | -0.230 | -0.005 (0.002) | **$4.3x10^{-3}$** | 82 | -0.005 | 0.000 (0.003) | $9.4x10^{-1}$ | 0.5171 |
| Skin - Sun Exposed | 286 | -0.260 | -0.007 (0.001) | **$9.6x10^{-7}$** | 187 | -0.220 | -0.006 (0.002) | **$1.0x10^{-3}$** | 99 | -0.352 | -0.008 (0.002) | **$4.6x10^{-4}$** | 0.2800 |
| Stomach | 420 | -0.367 | -0.008 (0.001) | **$7.6x10^{-9}$** | 271 | -0.413 | -0.009 (0.002) | **$2.6x10^{-7}$** | 149 | -0.285 | -0.004 (0.002) | **$4.8x10^{-2}$** | 0.1738 |
| Testis | | | | | 306 | -0.060 | -0.003 (0.002) | $1.1x10^{-1}$ | | | | | |
| Thyroid | 255 | 0.034 | 0.001 (0.001) | $6.3x10^{-1}$ | 144 | -0.042 | -0.001 (0.002) | $5.7x10^{-1}$ | 111 | 0.141 | 0.003 (0.002) | $1.4x10^{-1}$ | 0.1391 |
| Vagina | | | | | | | | | 108 | 0.058 | 0.002 (0.002) | $2.8x10^{-1}$ | |
| Whole Blood | 637 | -0.348 | -0.005 (0.001) | **$1.0x10^{-13}$** | 417 | -0.352 | -0.005 (0.001) | **$1.5x10^{-8}$** | 220 | -0.339 | -0.005 (0.001) | **$3.8x10^{-6}$** | 0.9984 |

**Table S5**

RTL and association with age within tissue types ($n \geq 75$ samples). Analyses were also stratified by sex. Linear mixed models were adjusted for age, sex (when appropriate), race/ethnicity, BMI, and donor ischemic time and technical factors (DNA concentration and sample plate) as a random effect. Interaction was analyzed between sex and age. *P*-values were obtained from likelihood ratio tests comparing model with and without age covariate or interaction (depending on model). Estimates with *p*-values <0.05 are shown in bold.

| | | | | | | | Association with LTL from ENGAGE | | | |
|---|---|---|---|---|---|---|---|---|---|---|
| SNP | Chromosome | Position | Gene | Closest Genes | Long Allele | Long Allele Frequency | β | se | Basepairs | p |
| rs11125529 | 2 | 54248729 | ACYP2 | ACYP2, TSPYL6, C2orf73, SPTBN1, PSME4 | A | 0.14 | 0.056 | 0.0101 | 66.9 | $4.5x10^{-8}$ |
| rs10936599 | 3 | 169774313 | MYNN | MYNN, ACTRT3, TERC, LRRC34, LRRIQ4 | C | 0.75 | 0.097 | 0.0083 | 117.3 | $2.5x10^{-31}$ |
| rs2736100 | 5 | 1286401 | TERT | TERT, MIR4457, CLPTM1L, SLC6A18, SLC6A19 | C | 0.51 | 0.078 | 0.0087 | 94.2 | $4.4x10^{-19}$ |
| rs7675998 | 4 | 163086668 | intergenic | MIR4454, NAF1, NPY1R, NPY5R, TKTL2 | G | 0.78 | 0.074 | 0.0091 | 89.7 | $4.3x10^{-16}$ |
| rs9420907 | 10 | 103916707 | STN1 (OBFC1 [hg19]) | STN1, SLK, SH3PXD2A, COL17A1, MIR936 | C | 0.13 | 0.069 | 0.0105 | 82.8 | $6.9x10^{-11}$ |
| rs2535913 | 14 | 72948525 | DCAF4 | DCAF4, ZFYVE1, DPF3, RBM25, PSEN1 | G | 0.66 | 0.035 | 0.0075 | 41.8 | $4.4x10^{-6}$ |
| rs3027234 | 17 | 8232774 | CTC1 | CTC1, LINC00324, PFAS, AURKB, BORCS6 | C | 0.79 | 0.021 | 0.0091 | 25.2 | $2.1x10^{-2}$ |
| rs8105767 | 19 | 22032639 | intergenic | ZNF208, ZNF257, ZNF676, ZNF43, ZNF100 | G | 0.29 | 0.048 | 0.0078 | 57.6 | $1.1x10^{-9}$ |
| rs755017 | 20 | 63790269 | ZBTB46 | ZBTB46, SLC2A4RG, LIME1, ZGPAT, ABHD16B | G | 0.13 | 0.062 | 0.0106 | 74.1 | $6.7x10^{-9}$ |

**Table S6**

Description of SNPs used to compute a polygenic SNP score reflecting longer telomere length (TL). Effects reported in terms of long allele and extracted from results from GWAS of leukocyte TL from ENGAGE consortium (Codd et al 2013). Top closest genes to SNP based on NCBI Refseq within 1 megabase window.

| Tissue | n | Correlation | $p_{correlation}$ | $\beta_{TLscore}$ (se) | $p_{TLscore}$ |
|---|---|---|---|---|---|
| Brain - Cerebellum | 189 | 0.170 | 0.0195 | 0.413 (0.191) | 0.0297 |
| Brain - Hippocampus | 135 | 0.123 | 0.1547 | 0.25 (0.199) | 0.1948 |
| Colon - Sigmoid | 162 | 0.086 | 0.2752 | 0.253 (0.272) | 0.3497 |
| Colon - Transverse | 483 | 0.111 | 0.0144 | 0.311 (0.132) | 0.0177 |
| Esophagus - GJ | 231 | -0.020 | 0.7576 | -0.182 (0.190) | 0.3267 |
| Esophagus - Mucosa | 466 | 0.030 | 0.5120 | -0.056 (0.123) | 0.6471 |
| Kidney - Cortex | 175 | 0.114 | 0.1331 | 0.263 (0.254) | 0.2853 |
| Lung | 473 | 0.057 | 0.2123 | 0.076 (0.093) | 0.4096 |
| Nerve - Tibial | 140 | 0.121 | 0.1550 | 0.236 (0.173) | 0.1588 |
| Ovary | 133 | 0.084 | 0.3390 | 0.273 (0.267) | 0.2903 |
| Pancreas | 459 | 0.104 | 0.0264 | 0.292 (0.140) | 0.0363 |
| Prostate | 162 | 0.162 | 0.0399 | 0.417 (0.221) | 0.0536 |
| Skin - Exposed | 255 | 0.046 | 0.4602 | 0.066 (0.180) | 0.7109 |
| Skin - Unexposed | 230 | 0.094 | 0.1555 | 0.129 (0.172) | 0.4425 |
| Stomach | 367 | 0.044 | 0.4041 | 0.033 (0.159) | 0.8345 |
| Testis | 269 | 0.029 | 0.6314 | 0.096 (0.275) | 0.7204 |
| Thyroid | 222 | 0.113 | 0.0930 | 0.161 (0.176) | 0.3479 |
| Whole Blood | 554 | 0.152 | 0.0003 | 0.200 (0.075) | 0.0073 |
| All Tissues | 5530 | 0.063 | $2.4\times10^{-6}$ | 0.133 (0.053) | 0.0113 |

**Table S7**

Association between a polygenic SNP score for long TL and measured RTL within and across tissues. Numbers of samples of each tissue were based on the number of tissues with both RTL and genetic sequence information available ($n \geq 100$). Pearson's correlation between RTL and the SNP score and corresponding *p*-value were reported. Linear mixed models were used to estimate the association (β) between ancestry and RTL within each tissue type and obtained *p*-values from likelihood ratio test comparing model with and without SNP score. For within tissue analyses, adjusted linear mixed models included sex, age, donor ischemic time, top five genotyping PCs, and technical factors (DNA concentration and sample plate) as a random effect. For across all tissue types analysis, adjusted linear mixed models included sex, age, donor ischemic time, top five genotyping PCs, technical factors, and the random effects of tissue type and donor.

| *TERT* Expression | | | | | Pearson Correlation | | | | Continuous (adjusted) | | Binary (adjusted) | |
|---|---|---|---|---|---|---|---|---|---|---|---|---|
| Tissue | n | Expressed | Not Expressed | mean (sd) | Age | p | RTL | p | β (se) | p | β (se) | p |
| Colon - Sigmoid | 101 | 26 | 75 | 0.146 (0.341) | -0.172 | 0.0848 | 0.104 | 0.3147 | 0.076 (0.098) | 0.4213 | -0.060 (0.081) | 0.4437 |
| Colon - Transverse | 299 | 229 | 70 | 0.588 (0.558) | -0.264 | **0.0000** | 0.109 | 0.0721 | 0.046 (0.033) | 0.1583 | 0.084 (0.044) | 0.0576 |
| Esophagus - Mucosa | 375 | 227 | 148 | 0.177 (0.171) | -0.100 | 0.0536 | 0.056 | 0.3012 | 0.036 (0.089) | 0.6782 | -0.021 (0.030) | 0.4923 |
| Lung | 449 | 43 | 406 | 0.033 (0.072) | 0.036 | 0.4457 | 0.031 | 0.5458 | -0.009 (0.178) | 0.9584 | 0.029 (0.038) | 0.4380 |
| Prostate | 88 | 8 | 80 | 0.033 (0.062) | 0.150 | 0.1640 | 0.058 | 0.5995 | 0.467 (0.558) | 0.3833 | 0.112 (0.124) | 0.3471 |
| Skin - Not Sun Exposed | 219 | 83 | 136 | 0.112 (0.131) | 0.138 | **0.0415** | -0.008 | 0.9098 | 0.072 (0.142) | 0.0151 | -0.014 (0.039) | 0.0233 |
| Skin - Sun Exposed | 278 | 97 | 181 | 0.112 (0.136) | 0.011 | 0.8571 | 0.162 | **0.0177** | 0.346 (0.144) | 0.6057 | 0.094 (0.042) | 0.7090 |
| Stomach | 205 | 117 | 88 | 0.192 (0.252) | -0.166 | **0.0171** | -0.025 | 0.7281 | -0.100 (0.092) | 0.2637 | -0.028 (0.049) | 0.5644 |
| Testis | 217 | 217 | 0 | 2.292 (0.837) | 0.007 | 0.9206 | -0.036 | 0.6164 | -0.021 (0.039) | 0.5789 | | |
| Thyroid | 199 | 15 | 184 | 0.038 (0.155) | -0.079 | 0.2650 | -0.046 | 0.5302 | -0.076 (0.125) | 0.5302 | 0.012 (0.072) | 0.8737 |
| Vagina | 75 | 12 | 63 | 0.036 (0.053) | 0.046 | 0.6923 | -0.004 | 0.9756 | -0.176 (0.714) | 0.7942 | -0.056 (0.086) | 0.4912 |
| Whole Blood | 496 | 53 | 443 | 0.043 (0.085) | -0.088 | 0.0511 | 0.088 | **0.0489** | 0.108 (0.111) | 0.3249 | 0.015 (0.030) | 0.5848 |

**Table S8**

RTL association with *TERT* gene expression. Expression was modeled as both a binary variable comparing expression (TPM $> 0.1$) to non-expression (TPM $\leq 0.1$) and as TPM (continuous). Tissues were analyzed if expression was detected in 5% or more of samples within that tissue type. Adjusted models included age, sex, race/ethnicity, BMI, donor ischemic time, and technical factors (DNA concentration and sample plate) as a random effect and corresponding *p*-values obtained from likelihood ratio test.

| *TERC* Expression | | | | | Pearson Correlation | | | | Continuous (adjusted) | | Binary (adjusted) | |
|---|---|---|---|---|---|---|---|---|---|---|---|---|
| Tissue | n | Expressed | Not Expressed | mean (sd) | Age | p | RTL | p | β (se) | p | β (se) | p |
| Brain - Cerebellum | 154 | 57 | 97 | 0.135 (0.203) | 0.099 | 0.2223 | -0.016 | 0.8459 | -0.055 (0.101) | 0.5929 | -0.035 (0.042) | 0.4075 |
| Brain - Hippocampus | 107 | 24 | 83 | 0.059 (0.120) | -0.079 | 0.4164 | 0.108 | 0.3064 | 0.157 (0.214) | 0.4461 | 0.019 (0.061) | 0.7407 |
| Colon - Sigmoid | 101 | 18 | 83 | 0.053 (0.134) | -0.107 | 0.2862 | 0.058 | 0.5786 | 0.040 (0.247) | 0.8680 | 0.082 (0.087) | 0.3270 |
| Colon - Transverse | 299 | 74 | 225 | 0.104 (0.222) | -0.121 | **0.0369** | 0.035 | 0.5630 | 0.026 (0.083) | 0.7484 | 0.024 (0.042) | 0.5544 |
| Esophagus - GJ | 156 | 44 | 112 | 0.096 (0.182) | 0.038 | 0.6349 | -0.129 | 0.1203 | -0.189 (0.119) | 0.1043 | -0.085 (0.051) | 0.0865 |
| Esophagus - Mucosa | 375 | 68 | 307 | 0.055 (0.129) | -0.052 | 0.3146 | -0.100 | 0.0649 | -0.261 (0.114) | 0.0214 | -0.090 (0.038) | 0.0170 |
| Lung | 449 | 53 | 396 | 0.037 (0.186) | 0.025 | 0.5902 | -0.048 | 0.3459 | -0.028 (0.057) | 0.6223 | -0.014 (0.035) | 0.6822 |
| Nerve - Tibial | 133 | 21 | 112 | 0.049 (0.124) | 0.044 | 0.6125 | -0.004 | 0.9674 | 0.153 (0.153) | 0.2999 | 0.042 (0.051) | 0.3961 |
| Ovary | 126 | 18 | 108 | 0.037 (0.100) | -0.094 | 0.2962 | 0.015 | 0.8668 | 0.283 (0.283) | 0.3193 | -0.021 (0.083) | 0.8003 |
| Pancreas | 263 | 21 | 242 | 0.019 (0.066) | -0.091 | 0.1416 | 0.014 | 0.8265 | -0.037 (0.277) | 0.8957 | -0.029 (0.068) | 0.6708 |
| Prostate | 88 | 11 | 77 | 0.038 (0.109) | 0.118 | 0.2727 | -0.148 | 0.1750 | -0.174 (0.318) | 0.5676 | -0.010 (0.103) | 0.9169 |
| Skin - Not Sun Exposed | 219 | 34 | 185 | 0.045 (0.120) | -0.135 | **0.0454** | 0.025 | 0.7195 | -0.055 (0.165) | 0.3248 | -0.024 (0.052) | 0.2654 |
| Skin - Sun Exposed | 278 | 37 | 241 | 0.052 (0.147) | 0.102 | 0.0882 | 0.040 | 0.5573 | 0.131 (0.136) | 0.7346 | 0.065 (0.059) | 0.6462 |
| Stomach | 205 | 54 | 151 | 0.074 (0.145) | -0.158 | **0.0240** | 0.067 | 0.3585 | -0.003 (0.163) | 0.9834 | -0.040 (0.055) | 0.4557 |
| Testis | 217 | 181 | 36 | 0.552 (0.425) | -0.097 | 0.1552 | 0.009 | 0.8980 | 0.029 (0.077) | 0.6979 | 0.039 (0.084) | 0.6387 |
| Thyroid | 199 | 37 | 162 | 0.076 (0.192) | 0.032 | 0.6502 | -0.007 | 0.9198 | 0.018 (0.104) | 0.8650 | 0.046 (0.051) | 0.3594 |
| Vagina | 75 | 13 | 62 | 0.065 (0.148) | -0.182 | 0.1175 | 0.038 | 0.7543 | 0.083 (0.221) | 0.6971 | 0.021 (0.087) | 0.8065 |
| Whole Blood | 496 | 31 | 465 | 0.016 (0.068) | 0.000 | 0.9976 | -0.053 | 0.2356 | -0.120 (0.139) | 0.3853 | -0.025 (0.039) | 0.5108 |

**Table S9**

RTL association with *TERC* gene expression. Expression was modeled as a binary variable comparing expression (TPM > 0.1) to non-expression (TPM ≤ 0.1) and as TPM (continuous). Tissues were analyzed if expression was detected in 5% or more of samples within that tissue type. Adjusted models included age, sex, race/ethnicity, BMI, donor ischemic time, and technical factors (DNA concentration and sample plate) as a random effect and corresponding *p*-values obtained from likelihood ratio test.

| *DKC1* Expression | | | Pearson Correlation | | | | Continuous (adjusted) | |
|---|---|---|---|---|---|---|---|---|
| Tissue | n | mean (sd) | Age | p | RTL | p | β (se) | p |
| Brain - Cerebellum | 154 | 26.153 (6.289) | -0.151 | 0.0622 | -0.057 | 0.4897 | -0.003 (0.003) | 0.3682 |
| Brain - Hippocampus | 107 | 9.729 (4.28) | -0.061 | 0.5329 | 0.048 | 0.6473 | 0.001 (0.006) | 0.8806 |
| Colon - Sigmoid | 101 | 18.976 (6.044) | -0.341 | **0.0005** | 0.081 | 0.4347 | 0.001 (0.006) | 0.8820 |
| Colon - Transverse | 299 | 21.792 (6.717) | -0.165 | **0.0043** | 0.018 | 0.7723 | 0.000 (0.003) | 0.8852 |
| Esophagus - GJ | 156 | 17.917 (4.439) | 0.069 | 0.3907 | -0.030 | 0.7153 | -0.001 (0.005) | 0.8789 |
| Esophagus - Mucosa | 375 | 34.452 (11.519) | -0.174 | **0.0007** | 0.109 | **0.0438** | 0.002 (0.001) | 0.2083 |
| Lung | 449 | 24.742 (8.906) | -0.110 | **0.0193** | 0.110 | **0.0293** | 0.002 (0.001) | 0.2370 |
| Muscle - Skeletal | 69 | 16.373 (5.952) | 0.007 | 0.9544 | 0.182 | 0.1529 | 0.009 (0.007) | 0.2163 |
| Nerve - Tibial | 133 | 25.774 (7.276) | 0.075 | 0.3880 | 0.096 | 0.3186 | 0.003 (0.003) | 0.3224 |
| Ovary | 126 | 31.023 (5.583) | -0.242 | **0.0063** | 0.123 | 0.1805 | 0.006 (0.005) | 0.2470 |
| Pancreas | 263 | 15.769 (5.691) | -0.035 | 0.5757 | -0.017 | 0.7901 | 0.000 (0.003) | 0.8915 |
| Prostate | 88 | 25.855 (7.051) | -0.017 | 0.8772 | 0.106 | 0.3331 | 0.003 (0.005) | 0.5893 |
| Skin - Not Sun Exposed | 219 | 33.711 (9.982) | 0.112 | 0.0968 | -0.081 | 0.2474 | -0.001 (0.002) | 0.5903 |
| Skin - Sun Exposed | 278 | 31.676 (8.139) | 0.072 | 0.2289 | 0.005 | 0.9435 | 0.001 (0.003) | 0.6427 |
| Stomach | 205 | 16.338 (7.208) | -0.212 | **0.0023** | 0.199 | **0.0056** | 0.006 (0.003) | 0.0656 |
| Testis | 217 | 35.708 (10.064) | -0.132 | 0.0524 | -0.108 | 0.1285 | -0.005 (0.003) | 0.1100 |
| Thyroid | 199 | 32.613 (6.575) | -0.040 | 0.5764 | 0.087 | 0.2321 | 0.004 (0.003) | 0.2072 |
| Vagina | 75 | 31.569 (10.419) | -0.143 | 0.2195 | -0.090 | 0.4557 | -0.003 (0.003) | 0.3173 |
| Whole Blood | 496 | 4.703 (3.423) | -0.073 | 0.1049 | 0.090 | **0.0457** | 0.004 (0.003) | 0.1874 |

**Table S10**

RTL association with *DKC1* gene expression. Expression was modeled as TPM (continuous) since all tissues had detectable expression of *DKC1* (TPM > 0.01). Tissues were analyzed if expression was detected in 5% or more of samples within that tissue type. Adjusted models included age, sex, race/ethnicity, BMI, donor ischemic time, and technical factors (DNA concentration and sample plate) as a random effect and corresponding *p*-values obtained from likelihood ratio test.

| Tissue | Total Normal Cells | Total Stem Cells | Number of Divisions per Stem Cell (per year) | Proportion of Stem Cells |
|---|---|---|---|---|
| Brain - Cerebellum | $8.46x10^{10}$ | $1.35x10^{8}$ | 0 | 0.0016 |
| Breast | $6.80x10^{11}$ | $8.70x10^{9}$ | 4.3 | 0.0126 |
| Prostate | $3.00x10^{10}$ | $2.10\ x10^{8}$ | 3.0 | 0.0070 |
| Lung | $4.34x10^{11}$ | $1.22x10^{9}$ | 0.07 | 0.0028 |
| Esophagus - Mucosa | $3.24x10^{9}$ | $9.98\ x10^{7}$ | 33.2 | 0.0299 |
| Colon - Transverse | $3x10^{10}$ | $2x10^{8}$ | 73 | 0.0066 |
| Colon - Sigmoid | $3x10^{10}$ | $2x10^{8}$ | 73 | 0.0066 |
| Whole Blood | $3.00x10^{12}$ | $1.35\ x10^{8}$ | 12 | 0.0000 |
| Pancreas | $1.70x10^{11}$ | $4.25x10^{9}$ | 1 | 0.0244 |
| Thyroid | $1.10x10^{11}$ | $7.15x10^{7}$ | 0.09 | 0.0006 |
| Skin - Unexposed | $1.84x10^{11}$ | $9.62x10^{9}$ | 5.55 | 0.0497 |
| Skin - Exposed | $1.84x10^{11}$ | $9.62x10^{9}$ | 5.55 | 0.0497 |

**Table S11**
Tissue specific stem cell estimates extracted from Tomasetti and Vogelstein et al (2015 and 2017). Estimates for number of divisions per stem cell and proportion of stem cells were analyzed for their association with *TERT* expression and RTL.

**Data S1.**

Pairwise Pearson Correlations between RTL measures taken from different tissue types. Each cell contains the Pearson correlation, sample size and *p*-value for each tissue pair

**Data S2**.
Difference in RTL between tissue pairs (ΔRTL) and its association with age. Correlation is dependent on pair order and significant correlations suggest pair differences that change with age. Linear models adjusted for sex, race/ethnicity, BMI, and donor ischemic time.

**Data S3.**
Colocalization posterior probabilities for telomere length (TL) associated gene regions. For selected tissue-specific eQTLs, where the lead TL-associated SNP was associated with expression with a $-\log_{10}$(*p*-value) that was > 0.9 when divided by the p-value for the lead SNP of the cis-eQTL, we reported the posterior probabilities (PP) from the co-localization analyses using the summary statistics from the ENGAGE GWAS of leukocyte TL and GTEx v8 cis-eQTL results under three sets of co-localization parameters corresponding to 50%, 20%, and 5% of causal TL SNPs affecting gene expression (see **Methods**). PP reported correspond to no causal variant (CV) (H0), causal variant for expression only (H1), causal variant for TL only (H2), two distinct causal variants (H3), and a common causal variant (H4).

**Data S4.**
Mediation analyses of telomere length on age-associated gene expression in whole blood. We utilized RNAseq TMM normalized and standardized gene expression data from whole blood (all genes expressed on standard normal (0,1)) and corresponding covariate files (GTEx v8). We identified age-associated gene expression within tissues from linear models adjusted for age, sex, donor ischemic time, RNA batch, and genotyping PCs and reported association estimate and p-value for age. We conducted mediation for RTL at each age-associated gene (adjusted for same covariates) and reported the average causal mediation effect (ACME) estimate, ACME p-value, total effect estimate, total effect p-value, and proportion mediation (ACME effect/total effect).

**Data S5.**
Mediation analyses of telomere length on age-associated gene expression in lung. We utilized RNAseq TMM normalized and standardized gene expression data from lung (all genes expressed on standard normal (0,1)) and corresponding covariate files (GTEx v8). See caption for **Data S4** for analysis information.

**Data S6.**
Mediation analyses of telomere length on age-associated gene expression in esophagus - mucosa. We utilized RNAseq TMM normalized and standardized gene expression data from esophagus - mucosa (all genes expressed on standard normal (0,1)) and corresponding covariate files (GTEx v8). See caption for **Data S4** for analysis information.

**Data S7.**
Geneset enrichment results of age-associated genes mediated by telomere length in whole blood. We enrichment age-associated genes mediated by TL in KEGG pathways and GO-terms using the *goana* and *kegga* functions in *limma* and extracted all terms with $p < 0.05$ (from Fisher's exact test), three or more TL-mediating genes, and between 10-5000 genes in pathway.

**Data S8.**
Geneset enrichment results of age-associated genes mediated by telomere length in lung.
See caption for **Data S7** for analysis information.

**Data S9.**
Geneset enrichment results of age-associated genes mediated by telomere length in esophagus - mucosa. See caption for **Data S7** for analysis information.
